# Discretized representations in V1 predict suboptimal orientation discrimination

**DOI:** 10.1101/2022.05.13.491867

**Authors:** Julien Corbo, O. Batuhan Erkat, John P. McClure, Hussein Khdour, Pierre-Olivier Polack

## Abstract

Neuronal population activity in sensory cortices is the substrate for perceptual decisions. Yet, we still do not understand how neuronal information content in sensory cortices relates to behavioral reports. To reconcile neurometric and psychometric performance, we recorded the activity of V1 neurons in mice performing a Go/NoGo orientation discrimination task. We found that, around the discrimination threshold, V1 does not represent the orientation of the stimuli as canonically expected. Instead, it forms categorical representations characterized by a relocation of activity at task-relevant domains of the orientation representational space. The relative neuronal activity at those discrete domains accurately predicted the probabilities of the animals’ decisions. Our results thus suggest that the categorical integration of discretized feature representations from sensory cortices explains perceptual decisions.

## Introduction

Perception results from the detection and interpretation of the information provided by the sensory organs in the central nervous system. It is delimited positively by a threshold above which a stimulus can be detected (detection threshold), and a limit above which two different stimuli can be discriminated (resolving power). While the detection threshold is constrained by the signal-to-noise ratio of the encoded stimulus (1), the resolution is in theory determined by the limit below which the neuronal activity patterns representing the two stimuli cannot be distinguished (2, 3). Currently, our models fail at predicting perceptual resolving power from neuronal activity (3, 4). Indeed, decoders that optimally utilize the information available in a neuronal population outperform the discrimination capability of the animals (5, 6, 7, 8, 4). Hence, a simple linear decoder using the neuronal activity of the primary visual cortex (V1) can distinguish the orientations of visual stimuli less than a degree apart (4), when mice cannot discriminate between orientations separated by an angle smaller than 20° (9, 10). Yet, it is well established that V1 neurons encode the orientation of visual stimuli (11, 12, 13), and V1 is necessary for resolving orientation discrimination tasks (6, 14, 15, 16, 17, 18, 19). This discrepancy of several orders of magnitude between psychometric and neurometric curves suggests that perceptual discrimination does not result from the neuronal information selection and integration approaches typically used by ideal observers. In that sense, the V1 readout realized by the system is suboptimal (4). It is therefore necessary to determine what properties of the V1 activity are relevant for perceptual decisions and how they constrain the discrimination abilities of the animals. As learning an orientation discrimination task was shown to reshape orientation representations in V1 (20, 21, 16, 22, 23), and since abstractive methods that optimally use the available neuronal information fail to capture the relationship between the V1 output and the perceptual abilities of the animal, we resolved to investigate the existence of a relationship between the V1 feature representations (i.e. orientation representations) and behavioral performance.

## Results

### Perceptual discrimination and neuronal population information content

To relate orientation representation and discrimination performance, we trained 10 mice to perform a Go/NoGo task that consisted in licking in response to a Go visual cue (a 45°drifting grating) and withholding licking during the third second of the presentation of the NoGo cue (a 135°drifting grating; **Fig. 1a**). Once they reached a satisfactory performance using orthogonal cues (sensitivity D’>1.7 for 3 consecutive sessions; **Fig. 1b**), we assessed the mice’s orientation discrimination abilities in six consecutive daily recording sessions during which the angle between the orientations of the Go and NoGo cues was progressively reduced (Go/NoGo angle decreasing from 90° to 15°; **Fig. 1a,b**; see Methods). This approach allowed us to record the V1 neuronal activity associated with behavioral performances above or below the discrimination threshold while keeping the mice engaged in the task. The performance of mice steadily decreased when the angle between the Go and NoGo cue was lower than 30°. This decrease was driven by an increase in the false alarm (FA) rate (**Fig. 1b**). On the last day, when the Go/NoGo angle was 15°, the D’ was systematically under the best random D’ value (D’ = 1.7; see Methods), indicating that the mouse orientation discrimination threshold was located between 30°and 15°, a value comparable to the orientation discrimination thresholds measured in similar studies using rodents (9, 10). During each session, we used calcium imaging to record the activity of V1 L2/3 cortical neurons as orientation discrimination task performance was previously shown to rely on V1 neuronal activity (14, 16, 17). The fractional fluorescence (ΔF/F) was then deconvolved in Action Potential related Events (20, 24) (APrEs; **Fig. 1c,d**). To confirm that the information available in V1 is sufficient to outperform the mice’s task performance, we trained shallow neuronal networks (SNNs) to classify trials as ‘Go’ or ‘NoGo’ using the V1 neuronal activity evoked by the task cues (**Fig. 1e**). At every Go/NoGo angle, the average classification error rate was below 2% (in descending order of Go/NoGo angle, mean±std : 0*±*5*10^*−*4^%, 0*±*4*10^*−*4^%, 0*±*4*10^*−*3^%, 0.2*±*0.5%, 0.3*±*1%, 1.5*±*2%), while it was 48.8*±*3% on average when the training set trials labels were shuffled (**Fig. 1f**). Therefore, SNNs vastly outperformed the animals (SNN D’ vs. mouse D’, mean±std: 4.65*±*0.009 vs. 3.0*±*0.39; 4.65*±*0.009 vs. 2.73*±*0.93; 4.65*±*0.07 vs. 2.06*±*0.76; 4.63*±*0.13 vs. 2.16*±*0.78; 4.61*±*0.19 vs. 1.41*±*0.64; 4.34*±*0.45 vs. 0.79*±*0.49; **Fig. 1g**). The information allowing such discriminability could be found both in pseudo populations using randomly chosen neurons across session, and in real populations using neurons recorded together during the same session (**Fig. 1h-j**). This result proves that the information present in V1 could theoretically allow the mice to classify the stimuli with a far better accuracy than their actual discrimination threshold (4). This discrepancy therefore suggested that the information present in V1 is not used to perform a pure semantic judgement (i.e., “Go or NoGo?”). Instead, the judgment must be based on feature information (i.e., the orientation of the cues). Indeed, orientation encoding is a core function of V1, inscribed in the network connectivity (25, 26, 27, 28), and is robust against the variation of other features (29, 30, 31, 32) and behavior (33). Therefore, we decided to test the hypothesis that constraints imposed by feature encoding in V1 could predict the suboptimal discrimination performance of the mice.

**Figure 1.**
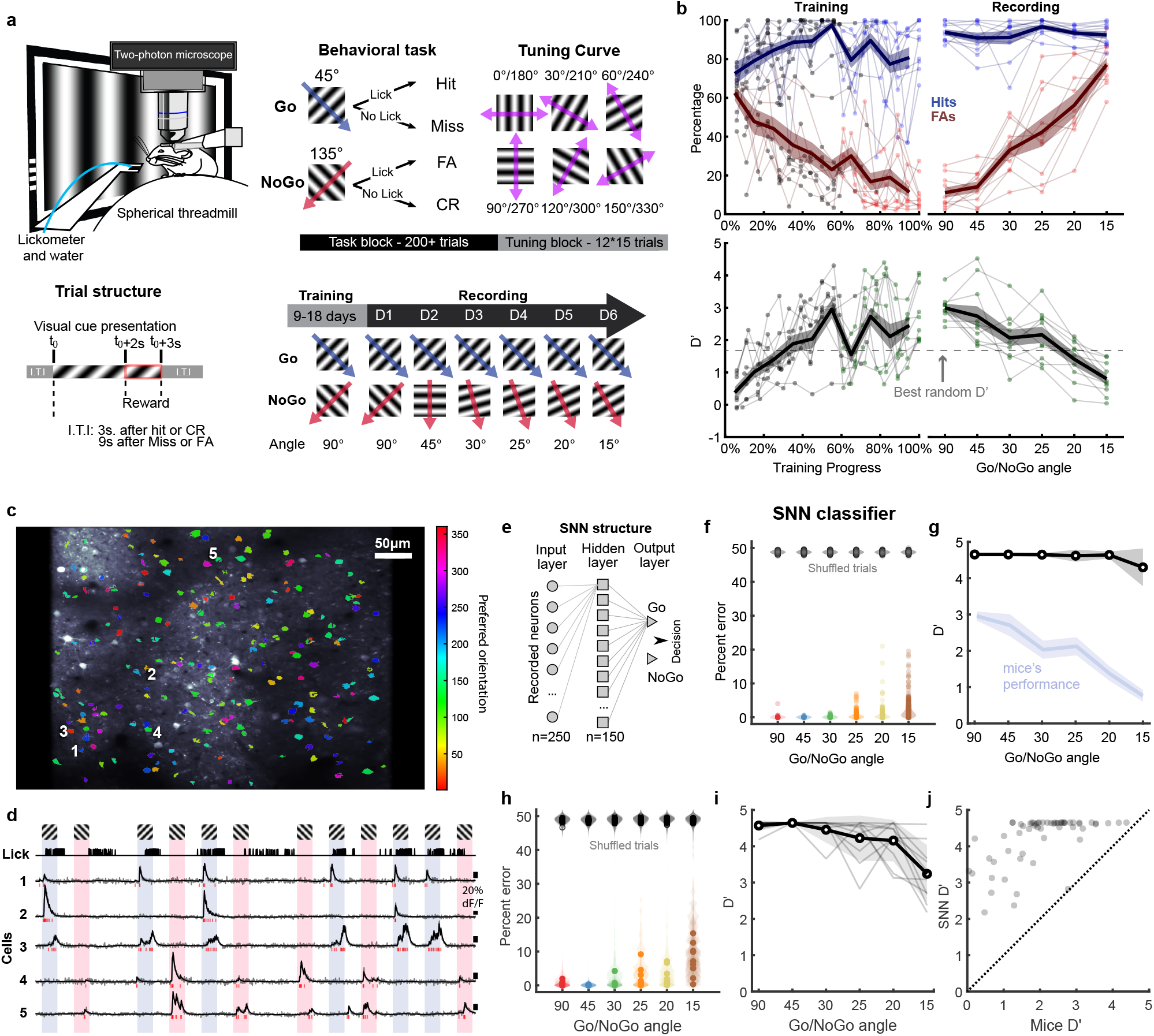
V1 neuronal activity contains enough information to outperform the animals in a discrimination task. **a**. Schematic representation of the experimental setup including the trial temporal structure (bottom left panel), the session organization (top right panel), and the Go and NoGo stimuli used for the different recording days (bottom right panel) **b**. Mice behavioral performance during training and recording sessions. Top panels: chance of hit trials (lick) during the presentation of the Go cue (blue lines), and of false alarms (FAs), i.e. licking during the presentation of the NoGo cue (red lines). **c**. Example of a two-photon calcium image on which were overlapped the neurons’ preferred orientations. **d**. Fractional fluorescence (ΔF/F, bottom traces) of the five selected neurons indicated in (c.) during the orientation discrimination task. Neurons 1-3 are tuned to the Go orientation. Neurons 4 and 5 are tuned to the NoGo orientation. Top trace: mouse licking activity. Short red lines indicate time of action potential related events (APrEs) provided by deconvolution. **e**. Schematic of the shallow neural network (SNN) used to classify the Go and NoGo cues based on the V1 evoked activity. **f**. Percent of correct classification of SNNs trained, using randomly selected neurons forming pseudo sessions. Circles: performance of the 1,000 trained SNNs; violin plots: performance distribution. Colors show the different Go/NoGo angles. Black datapoints: performance of SNNs trained with shuffled labels. **g**. D’ derived from the SNN performance shown in (f.) (black) compared to the mice behavioral performance (blue). The shaded grey area indicates the standard deviation. **h**. Same as (f.) but SNNs were trained with data provided by single recording sessions. Each circle indicates the average of the 100 repetitions of the procedure for each recording session; violin plots: performance distribution for the 100 repeats of each session. **i**. Same as (g.) but using the data from (h.). **j**. Scatter plot of the mice’s individual behavioral performance against the classification performance of the SNNs using their recorded neurons.

### Discrete domains in the orientation space

To understand how the orientation of task stimuli is encoded by the V1 neuronal population, we sorted V1 neurons by their preferred orientation (**Fig. 2a**), as the tuning properties of V1 neurons defines a feature encoding space for orientation (34, 32, 35, 36, 37, 4, 38, 39). Projecting the neuronal activity in that preferred orientation space allows therefore to access the orientation-related information in the V1 population responses. The preferred orientation of each neuron was assessed during a tuning block, i.e., a set of drifting gratings of 12 different orientations passively viewed by the animals (**Fig. 1a**). Tuning blocks were typically presented at the end of the recording sessions but presenting them at the beginning of the recording led to similar estimate of the neurons’ orientation preference (**Fig. S1**). When the Go/NoGo angle was higher than or equal to 45°(at the best task performance), the representations of the Go and the NoGo cues in the orientation space were completely distinct (**Fig. 2a, b; S2**), and those activity profiles were centered on the presented orientations. For Go/NoGo angles ranging from 30° to 20° (i.e., around the behavioral discrimination threshold), the overlap in the orientation space between the responses evoked by the Go and NoGo cues started to increase (**Fig. 2cf; S2**). Concurrently, the V1 responses in the orientation space to the NoGo cues stopped being a unimodal distribution centered around the orientation of the presented stimulus (**Fig. 2c, d; S2**). Instead, the NoGo cue representations were bimodal with a peak around 45° (i.e., close to the orientation of the Go visual stimulus) and another that remained around 90°regardless of the orientation of the NoGo stimulus. The bimodality of the NoGo stimulus representation in the orientation space disappeared when the Go/NoGo angle was 15°(when task performance was at worst; **Fig. 2e, f**). The distribution of the peak of activity in the orientation space for resampled pseudo-trials similarly confirmed that for Go/NoGo angles between 45°to 20°, the neuronal activity evoked by the Go and NoGo cues activated two distinct ‘domains’ of the orientation space: a “Go domain” (around 45°) and a “NoGo domain” (around 90°) that did not correspond to the orientations of the presented NoGo stimuli (**Fig. 2g**). the Go representations were also found to be distorted, as they were systematically skewed away from the NoGo orientation (**Fig. S2g**). To determine if those domains existed in the V1 of naïve mice that were not trained to perform the orientation discrimination task, we recorded the same sequence of 6 recording days in naïve mice passively viewing the ‘Go’ and ‘NoGo’ stimuli (**Fig. S3a**). The representations of those stimuli were all unimodal and the activity evoked in the orientation space was centered around the task cue orientation (**Fig. S3b-h**). Hence, between a Go/NoGo angle of 30° to 20°, the activity profiles evoked in trained mice presented a systematic activation of two separate domains of the orientation space that did not exist in naïve animals (**Fig. 2h-i**). This bimodal representation was not an artefact due to averaging the activity across trials, sessions, animals and similarly tuned neurons (**Fig. S4a**). Indeed, the simultaneous activation of the Go and NoGo domains was consistently present at the single trial level when the Go/NoGo angle was between 30° and 20° (**Fig. S4b**). Finally, it was not due to an alteration in the distribution of preferred orientations in the V1 population (**Fig. S5a**) nor to a change in the number of neurons responsive to the task stimuli (**Fig. S5b**). Our results therefore show that the Go and NoGo stimuli activate two discrete domains of the orientation space during the task. We hypothesized that this discretization of the activity could be relevant for the perceptual decisions. To test the existence of a relationship between the relative amount of activation of those domains and the behavioral performance, we needed to be able to manipulate the performance or the representation of orientations in V1 while maintaining identical the cue orientations and all the other task parameters. That way, we could test whether a change in the orientation representation of the same stimuli underpinned a change in performance. The presence of sounds during the presentation of visual cues has been shown to improve orientation discrimination in humans (40, 41) as well as the representation of oriented stimuli in the mouse V1 (42, 43). Therefore, we trained a second set of mice but this time we used two types of blocks during the recording sessions: unimodal blocks where mice performed in the same condition as the first set of mice, and audiovisual blocks during which the same visual stimuli were randomly paired with a pure tone (10 kHz or 5 kHz; **Fig. 3a**). We found that sounds improved the mouse orientation discrimination performance within the same session (**Fig. 3b**). This perceptual improvement was driven by the decrease in the FA rate (**Fig. 3c; S6**). The maximal improvement was found when the Go/NoGo angle was 25° (**Fig. 3c**), i.e., when the mouse performance during the unimodal blocks was for the first time below the best D’ value achievable with random choice behavior (D’ = 1.0; **Fig. 3b**). We then determined the representations of the Go and NoGo cues in the V1 orientation space during the unimodal and audiovisual blocks (**Fig. 3d; S6**). The results reproduced our initial findings. For wide Go/NoGo angles and below the orientation discrimination threshold (i.e., Go/NoGo angles *≥* 45°or *≤* 20°), the Go and NoGo cues were represented in the orientation space by a unimodal distribution centered on the orientation of the presented stimulus. In the vicinity of the discrimination threshold (*∼*25°), the representation of the NoGo cue was bimodal with a peak of activity around the orientation of the Go cue and another around 90°which did not correspond to the orientation of the NoGo stimulus (**Fig. 3d,e; S6**), reproducing the result obtained with our first dataset (**Fig 2**). To determine if the behavioral improvement found during the audiovisual blocks was associated with a modification of the relative activation of the Go and NoGo domains, we compared the representation of the NoGo orientation in the unimodal and audiovisual contexts. We found that the largest peak of activity in the unimodal context (mean D’ ± s.e.m. = 0.8±0.2; poor performance) was in the Go domain (45°). While, in the audiovisual context (D’ = 1.5±0.1; improved performance), the largest peak of activity was located in the NoGo domain (**Fig. 3d, e; S6**). Consequently, the relative activation of the Go and NoGo domains changed in the audiovisual context (sum of activity in the Go domain [1-60°, n=117 neurons] / sum of activity in the NoGo domain [61-120°, n=141 neurons], visual only vs. audiovisual, 1.43 vs. 1.13, permutation test p=1.45e-04, effect size Hedge’s g=0.23). Altogether our results demonstrated in two independent datasets that the task stimuli are represented in the orientation space by the activation of two neuronal populations: a population tuned for orientations surrounding the Go stimulus (the Go domain), and another population tuned for orientations surrounding 90° (the NoGo domain). The NoGo domain did not correspond to the orientation of the presented NoGo stimulus and remained similar across Go/NoGo angles and across datasets. Importantly, those findings could not be attributed to differences in locomotion or pupil size (**Fig. S7 and S8**).

**Figure 2.**
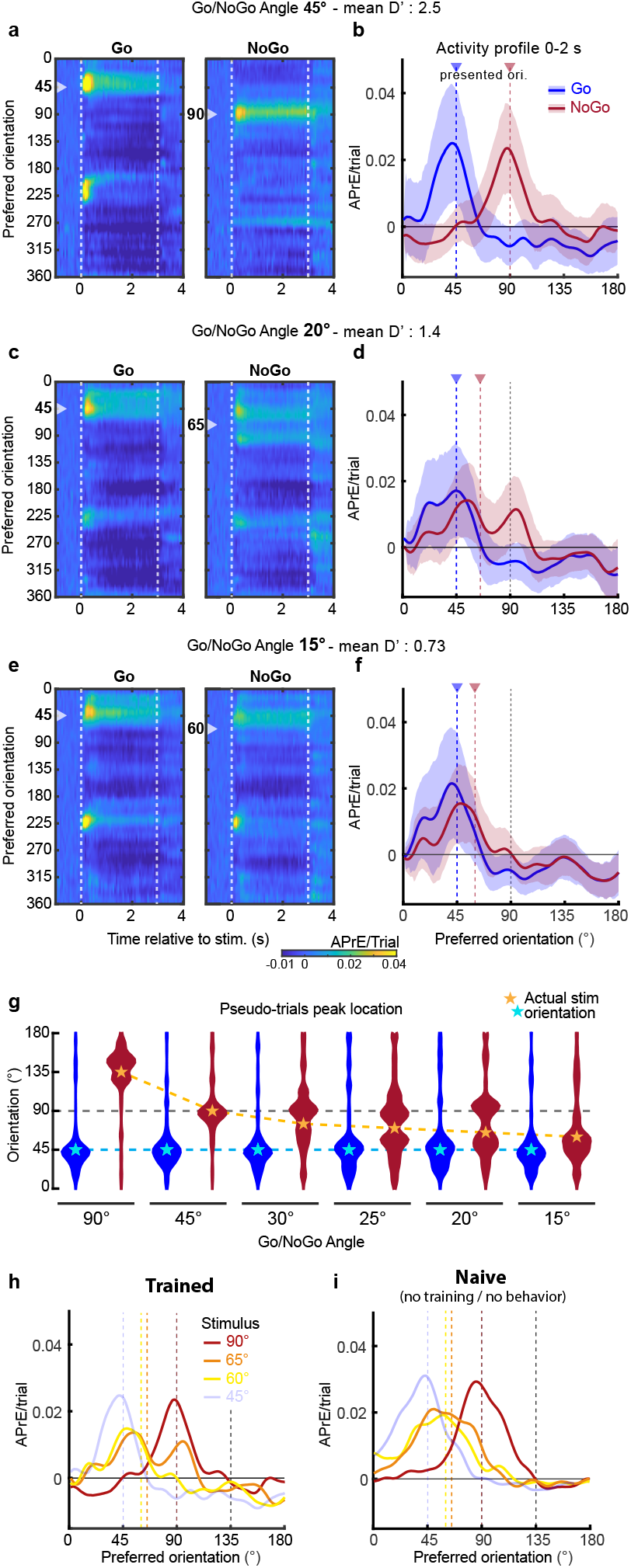
Discretized representations of the task cues in the V1 orientation space. **a**. Action Potential related Events (APrEs) generated by the presentation of the Go (45°, left panel) and NoGo stimuli (90°, right panel; Go/NoGo cue angle: 45°) as a function of the V1 neurons’ preferred orientation (average across all neurons of same preferred orientation). The neuronal activity color scale is located at the bottom of the panel (c). The mean behavioral performance (D’) across mice is indicated above the panel. **b**. Mean activity evoked in the orientation space by the Go (45°, blue) and NoGo (90°, red) stimuli during the two first seconds of the stimulus presentation (average between t=0s and t =2s of the activity shown in a.). Vertical colored lines indicate the orientation of the presented stimulus. Shaded areas indicate the s.e.m. **c**. Same representation as in (a) when the angle between the Go and NoGo cues is 20°. **d**. Same representation as in (b) when the Go orientation is 45°and NoGo orientation is 65°. **e**. Same representation as in (a) when the Go/NoGo angle is 15°. **f**. Same representation as in (b) when the Go orientation is 45°and the NoGo orientation is 60°. **g**. Violin plot of the distribution of the location of the peak of activity in the orientation space for Go (blue) and NoGo (red) pseudo-trials (n=1000). Blue and gold stars indicate the orientation of the Go and NoGo stimuli respectively. **h-i**. Average activity profiles evoked by the 45°, 90°, 65°and 60°stimuli in trained and naïve animals, respectively.

**Figure 3.**
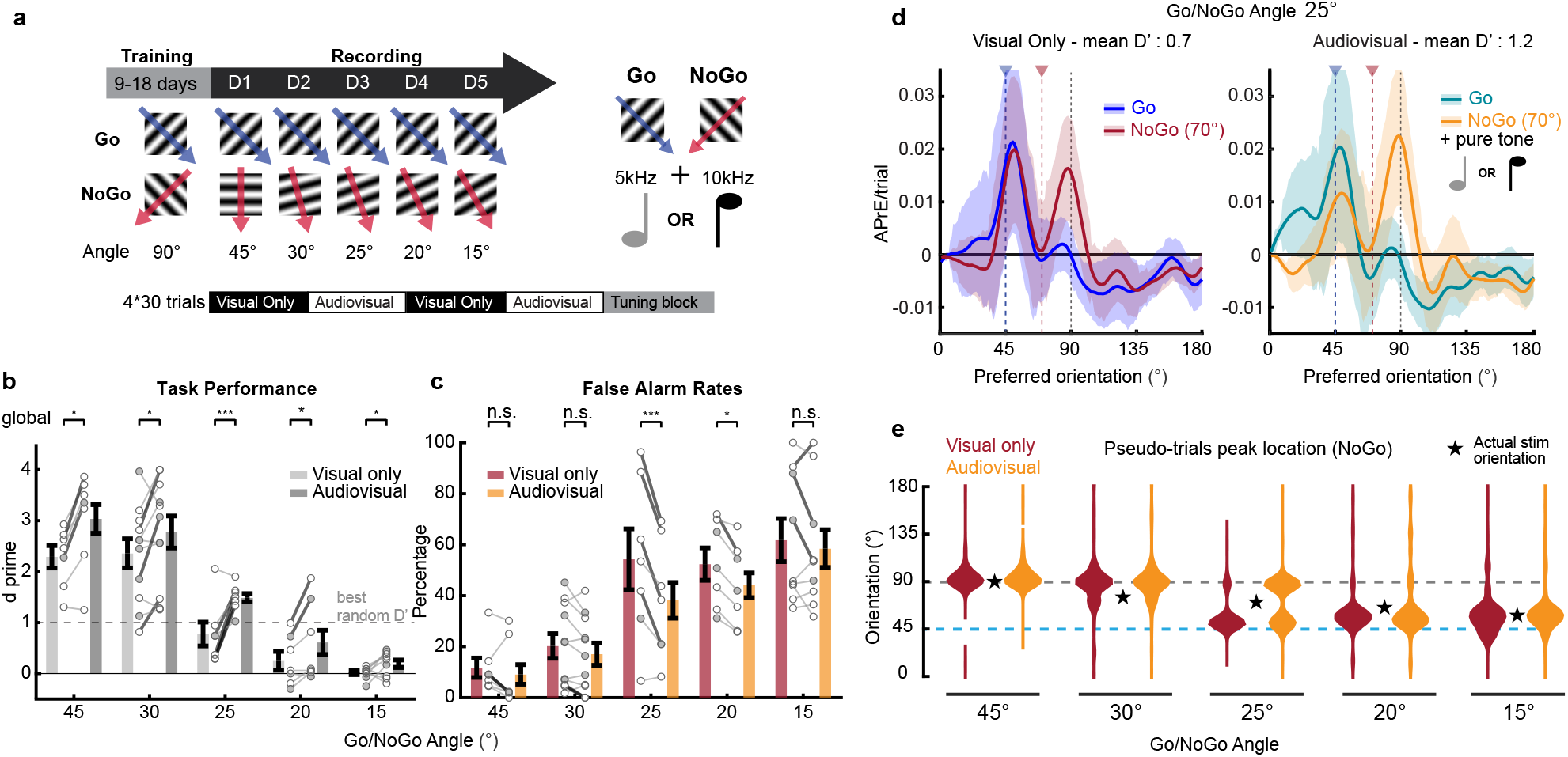
Orientation representation and behavioral performance in unimodal and audiovisual contexts. **a**. Go and NoGo stimuli used during the training and recording sessions (top left panel) for a second experimental group for which we alternated unimodal and audiovisual blocks. During audiovisual blocks Go and NoGo cues were presented simultaneously with either a 5 kHz or a 10 kHz tone. Recording sessions ended with a tuning block (bottom panel). **b**. Behavioral performance of the mice during the unimodal visual (light grey) and audiovisual (dark grey) blocks of the different recording sessions. Black lines indicate mean ± s.e.m. across sessions. Permutation test: *: p<0.05; ***: p<0.001. Circles indicate individual mice (thick lines indicate significant difference between unimodal and audiovisual performances; p<0.05). Grey circles indicate sessions for which only the behavior data was used. **c**. Same representation as in (b.) for the FA rate. **d**. Left panel: mean activity evoked in the orientation space by the Go (45°, blue) and NoGo (70°, red) cues during the two first seconds of the unimodal visual stimulus presentation. Right panel: mean activity evoked in the orientation space by the same Go (45°, green) and NoGo (70°, yellow) stimuli during the same period in the audiovisual context. Vertical colored lines indicate the orientation of the presented stimulus. Shaded areas indicate the s.e.m. **e**. Violin plot of the distribution of the location in the orientation space of the peak of activity for unimodal NoGo (red) and audiovisual NoGo (yellow) pseudo-trials. Black stars indicate the orientations of the presented NoGo stimuli.

### Categorical representations predict perceptual performance

The results obtained with the audiovisual dataset suggested that the relative activation of two neuronal populations in discrete domains of the orientation space was driving the animals’ behavioral performance. To test this hypothesis, we defined the Go evidence as the ratio between the APrE activity evoked in the Go domain (1-60°) and the sum of APrE activity in the Go and NoGo domains (61-120°; **Fig. 4a**). For each Go/NoGo angle of the two datasets, we plotted the mice’s licking probability as a function of the Go evidence for every time point of the visual stimulus presentation (example in **Fig. 4b** for t = 1.8-2s). The linear fit between the Go evidence and the mouse task performance explained more than 80% of the behavioral performance variance within the first 500 milliseconds of the stimulus presentation (**Fig. 4c**). The variance explained by the Go evidence was significantly greater than that explained by the Go domain activity or the NoGo domain activity alone (**Fig. 4c**). This suggested that neuronal activities of both domains were relevant to the animal’s decision. The relationship between Go evidence and behavior did not depend on specific parameters for the Go and NoGo domains (center or width) as long as the Go and NoGo domains contained 45° and 90° exclusively (**Fig. S9**). Therefore, we demonstrated that the relative activation of the neuronal populations of the Go and NoGo domains accurately predicts the behavioral choice probabilities. This relationship between V1 neuronal activity and perceptual behavior was initially found using the averaged neuronal responses including successful and unsuccessful trials. We therefore needed to test the hypothesis that CRs could result from the sole activation of the NoGo domain while FAs were the consequence of the sole activation of the Go domain. Reciprocally, Hits could result from the sole activation of the Go domain while Misses would be the consequence of a strong activation of the NoGo domain. However, this hypothesis was ruled out as the activity profiles evoked during FA and correct rejection (CR) trials overlapped in the orientation space (**Fig. 4e-f, S10**). Moreover, this invariance of the stimulus representations across behavioral decisions was also found at the single trial level (**Fig. 4g**). We plotted the activity evoked by the task stimuli in the Go vs. NoGo domains to determine if the single trials’ activity tended to cluster in distinct locations of the Go evidence space according to the decision (hit vs. miss, CR vs. FA; **Fig. 4h**). In 54% of the recordings the single trials’ clouds did not differ between successful and unsuccessful trials (2D Kolmogorov-Smirnoff test p>0.05, 25/60 for Go, 40/60 for NoGo). For the remaining sessions, we computed the translation of those clouds between successful and unsuccessful behavioral responses (**Fig. 4i**). The vectors were not isotropically oriented (**Fig. 4i-j**, Watson’s U test for anisotropy, p=0.013). When the directions of the displacement vectors were measured relatively to the local direction of isoevidence (**Fig. 4k**-top), i.e. the line along which the Go evidence value remains constant, their average angle was not different from 0 (**Fig. 4k**-bottom, ttest p=0.99). This means that when there was a difference in neuronal activity between successful and unsuccessful trials, this modulation was orthogonal to a modulation of the Go evidence, i.e., both Go and NoGo domain were similarly modulated. Such orthogonal modulation is likely to originate from the difference in movement and/or arousal during licking responses (**Fig S7**, Go trials are associated with more licking and more locomotion). As the orientation representation of the Go and NoGo cues were the same for successful and unsuccessful trials, V1 cannot be considered as the location where the perceptual decision emerges. Instead, it seems that the ratio between the neuronal activity evoked in the Go and NoGo domains represents a probability that the presented stimulus belongs to either the Go or NoGo category. Hence, choice variability likely arises outside of V1 (44, 4).

**Figure 4.**
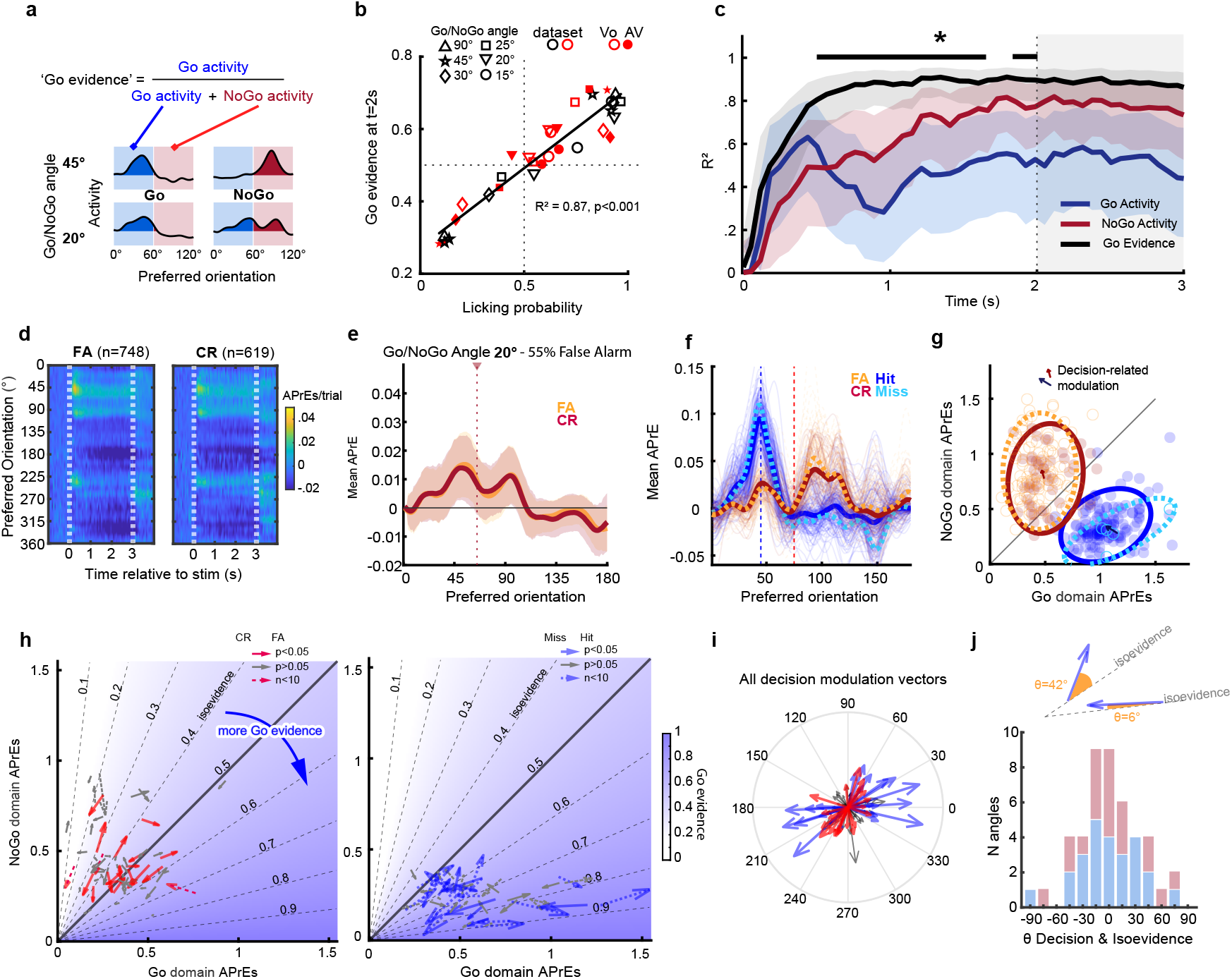
Categorical representations predict the mice’s behavioral performance. **a**. Delimitation of the Go (1°-60°, blue) and NoGo (61°-120°, red) domains in the orientation representational space. Go evidence is defined as the ratio between the amount of neuronal activity in the Go domain (area under the activation profile) and the sum of the neuronal activity in the Go and NoGo domains. **b**. Mouse licking response probability as a function of the Go evidence measured during the 200 ms preceding the beginning of the response window for all the sessions of the two datasets. For the second dataset (red), we separated unimodal (empty symbols) and audiovisual (solid symbols) blocks. Black line: linear fit (r2: 0.87; p<0.001). **c**. Value of the linear fit Pearson correlation coefficients measured as in (b) for all the timepoints during the time course of the visual stimulus presentation. Black line: Go evidence; blue line: Go domain activity; red line: NoGo domain activity. Shaded areas indicate the 95% confidence interval of the correlation coefficient. *: significant difference between the Pearson correlation coefficients measured using the Go evidence and using the Go and NoGo activity (significance is reached when both values are located outside of each other’s confidence interval). **d**. Activity evoked in the orientation space by the 65°NoGo stimulus (Go/NoGo angle 20°; visual-only dataset) in FA and CR trials. **e**. Average activity profiles in the orientation space of FA and CR trials during 0-2s of stimulus presentation using the data in d. **f**. Single trial and average activity profiles evoked by the Go and NoGo cues (angle 20°) for an example session. Trials are sorted by the animals’ decision. Dashed lines: average profile of the unsuccessful trials. **g**. Activity in the Go and NoGo domains as defined in (a.), for all the trials of the example session. The ellipses are fit to the clouds of single trial values. Arrows show the displacement of the centroid of the ellipses fitting the correct and incorrect decisions. **h**. Displacement of the centroid of the neuronal activity in the Go and NoGo domains associated with the perceptual decision for Go (Left) and NoGo (right) trials. Each vector corresponds to a session. The vectors point towards the centroids of trials in which the decision was to lick. Grey arrows indicate non-significant translations (two-dimension Kolmogorov-Smirnoff test). Dotted lines indicate sessions for which there were less than 10 trials with one of the decision types. The background color indicates the Go evidence (see a.) value at every location of the Go-NoGo domain activity space. **i**. Vectors showed in (h.) plotted from the origin of a polar space. **j**. Top: examples illustrating the measure of the angle between the line of isoevidence and the centroid translation vector. Bottom: distribution of angles between centroid translation and isoevidence, for Go (blue) and NoGo (red) stimuli.

### A modulation of neuronal excitability underpins the discretization of task orientation representations

In a previous study, we showed that distortions of the orientation representation in V1 could result from local suppressions of the neuronal excitability in the orientation space (20), as measured by the amplitude of the response to the neurons’ preferred orientation. Therefore, we estimated the neuronal excitability in the orientation space by measuring the amplitude of the tuning curves for all the oriented V1 neurons (**Fig. 5a**). In trained animals, neurons tuned to 45±7.5° and 90±7.5° had a similar peak amplitude (mean ± s.e.m, respectively 1.8±0.2 and 1.5±0.1 APrEs, permutation test p = 0.15, hedge’s g for effect size = 0.12). However, neurons tuned for 67±7.5° (at the location where the trough of activity separating the Go and NoGo domains) had a lower tuning curve amplitude (1.0±0.1 APrEs; amplitude vs. 45° neurons’ group: p=0.0001, g=0.43; amplitude vs. 90°neurons’ group: p=0.0002, g=0.41). To continuously assess the tuning curve amplitudes in the orientation space, we grouped the neurons in 7.5°bins, then plotted their mean tuning curve amplitude as a function of their orientation preference (**Fig. 5b**). For every Go/NoGo angle, we found the presence of the same W-shaped modulation of the neuronal excitability in the orientation space. This modulation was characterized by two troughs of low excitability (mean location and 95% confidence interval: 68.3°, [66-70°] and 105.8°, [105-108°]); **Fig. 5c**) that corresponded to the flanks of the Go and NoGo orientations used during training, separated by a return to baseline excitability peaking at 88.3° (95% confidence interval: [86-91°]). The same analysis performed on data from naïve mice demonstrated the absence of such modulation of the neuronal excitability in the orientation space (**Fig. 5d**, average number of APrEs in the same preference intervals and 95% confidence interval: first through [66-70°]: 1.17 [1.11-1.23]; second through [105-108°], 1.29 [1.22-1.37]; center peak [86-91°]: 1.28 [1.22-1.34]). We then tested the hypothesis that the W-shaped modulation of neuronal excitability in the orientation space could explain the distorted representations of the task stimuli. Using a qualitative model, we showed that the modulation of the neuronal excitability in the orientation space can transform an orientation-unbiased input (a gaussian distribution of the neuronal activity centered in the orientation space on the orientation of the Go or the NoGo cue) into the profiles obtained experimentally for all the Go/NoGo angles (**Fig. S11a**,**b**). Moreover, pure tones had previously been shown to sharpen orientation representations in the orientation space (43, 45). Hence, we tested the effect of the neuronal excitability modulation on sharper orientation inputs. We found that the resulting activity profiles evoked by the Go and NoGo cues at all Go/NoGo angles reproduced the experimental results obtained in the audiovisual blocks of our second dataset, i.e., a larger NoGo signal at the behavioral threshold in the presence of sound (**Fig. S11c**). Altogether, this qualitative analysis suggests that the bimodal representation of the NoGo cues around the discrimination threshold can be explained by a simple multiplicative interaction between orientation-unbiased inputs and a local decrease in neuronal excitability around the task values.

**Figure 5.**
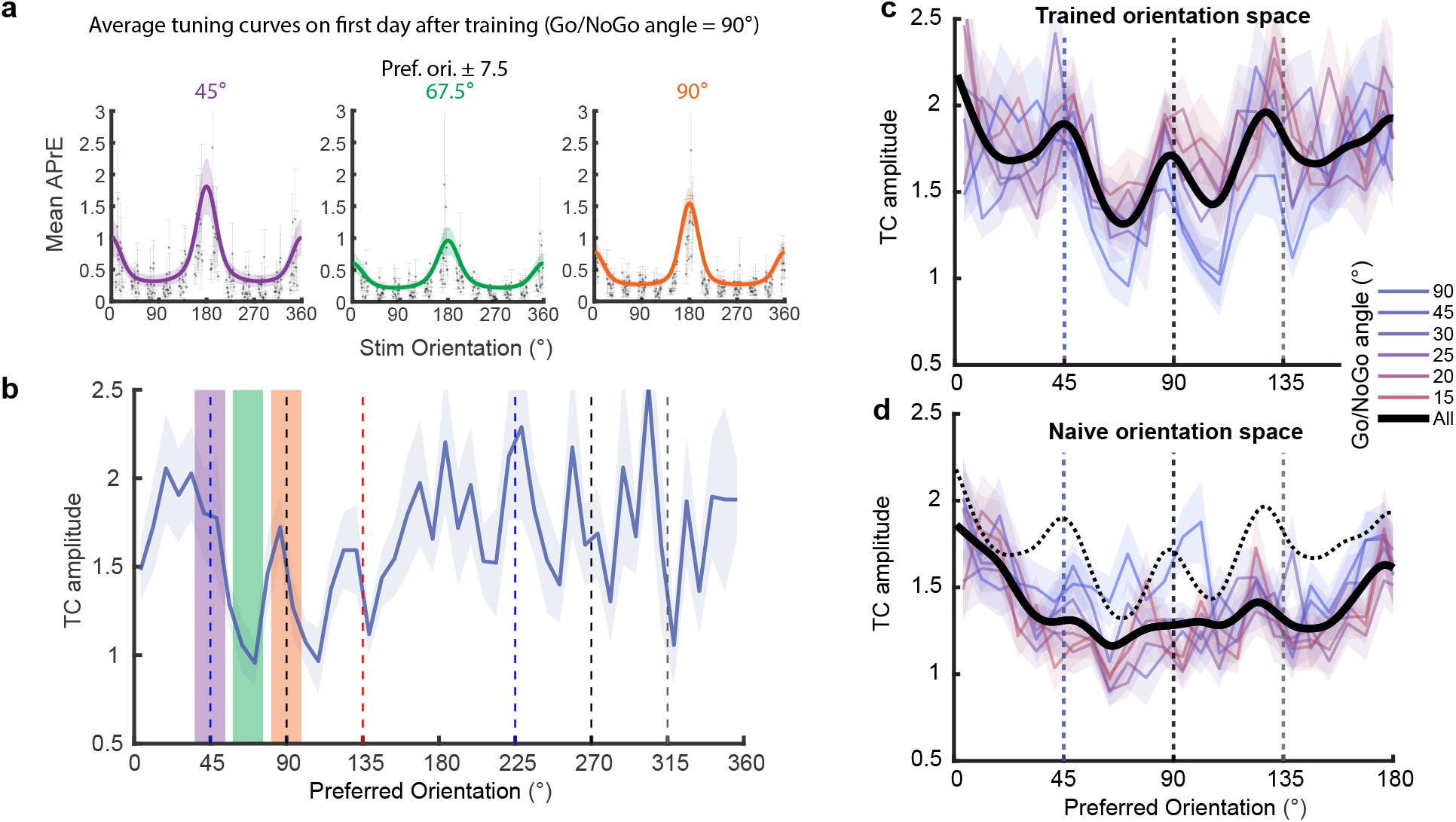
Modulation of the neuronal excitability in the orientation space. **a**. Tuning curves of the neurons tuned for 45 ±7.5°(left), 67.5 ±7.5°(center), and 90 ±7.5°(right), recorded during the 90°Go/NoGo angle sessions. Grey dots indicate mean activity, grey vertical lines indicate s.e.m. across neurons. Colored lines and shadow indicate mean tuning curve fit ± s.e.m. across neurons. **b**. Mean amplitude of the tuning curve as a function of the neurons’ preferred orientation for the sessions when the Go/NoGo angle was 90°(shadow indicates s.e.m. across neurons of similar preferred orientation). **c**. Modulation of the neuronal excitability in the orientation space for the sessions with all the different Go/NoGo angles (color code on the right). Black trace: averaged modulation across Go/NoGo angles, fitted with a smoothing spline. **d**. Same as c. but for the naïve animals. The dashed line shows the average from (c).

### Relationship between representational similarity and behavioral performance

Recent studies having suggested that discriminability results from the activation of distinct neuronal populations (46, 47, 48), we hypothesized that the variable amount of overlap in the discrete orientation domains should translate into correlated amounts of representational similarity. To test this hypothesis, we measured for each recording session the cosine similarity between the responses evoked by the task cues (**Fig. 6a-c**). We found a significant relationship between the animals’ D’ and the similarity of task stimuli representations (**Fig. 6d**, R2 = 0.43, F-test against constant model F=12.06, p=4.3*10-5, inflection point: similarity=0.73, D’=1.37). We then looked for the minimum level of similarity necessary to discriminate between two orientations. We first computed the selfsimilarity of Go and NoGo representations (i.e., the similarity across trials of the same stimulus, **Fig. 6e-f**). This value was stable across recording days (for the Go cue: One-way Anova, F(5,54)=0.19, p=0.96, for the NoGo cue: F(5,54)=0.28, p=0.92). Then, we showed that the self-similarity was larger than the similarity between the Go and NoGo responses, except when the angle between the Go and NoGo cues was 15°(permutation test, p<0.004 for all Go/NoGo angles except for 15°; Go/NoGo similarity at 15°against Go self-similarity, mean±std: 0.78±0.03 vs 0.81±0.07, p=0.21; against NoGo self-similarity: 0.78±0.03 vs 0.79±0.05, p=0.57). This corresponded to the angle at which animals performed below the chance threshold (**Fig. 1b**). Hence, we confirmed that an excess of similarity in the V1 population responses is detrimental for perceptual discrimination.

**Figure 6.**
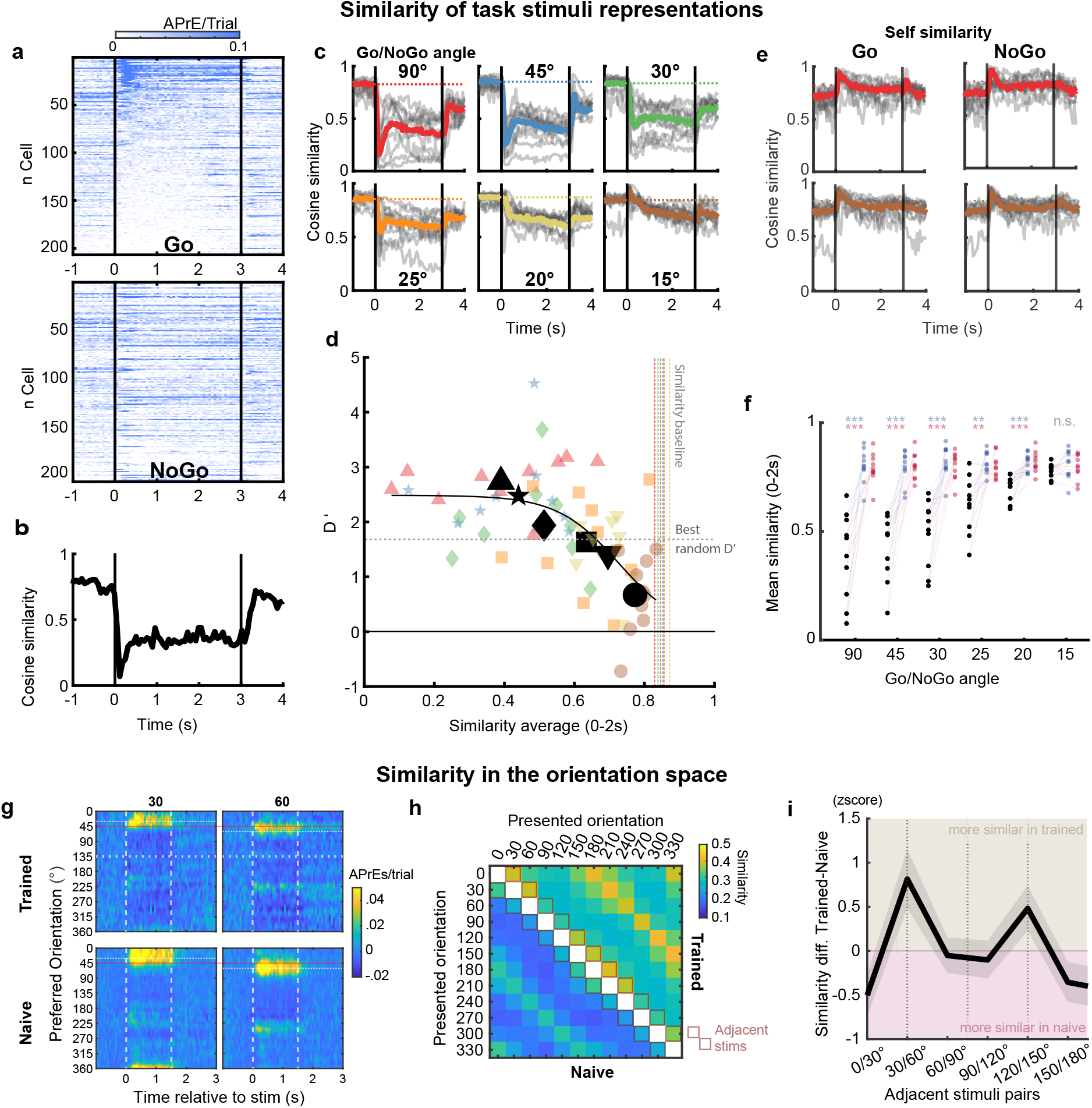
Cosine similarity between the representations of the task and tuning block stimuli. **a**. Trial-averaged APrE activity of 206 neurons simultaneously recorded in an example session (Go/NoGo angle = 90°), during the presentation of the Go (top panel) and NoGo cues (bottom panel). In both panels, neurons are ordered as a function of their response intensity to the Go stimulus presentation. **b**. Cosine similarity between the neuronal activity patterns evoked by the Go and NoGo cues that are shown in (a). **c**. Same representation as in (b) for all the Go/NoGo angles of all the mice (thin grey lines: individual mice; colored lines: averages across mice). **d**. Mouse behavioral performance (D’) as a function of the average cosine similarity between the Go and NoGo neuronal responses during the first two seconds of the visual stimulus presentation. Color code for individual sessions correspond to the one used in (c). Black shapes indicate the mean across mice for the different Go/NoGo angles. Vertical colored lines indicate the cosine similarity measured during the baseline activity immediately preceding the visual stimulus (duration: 1s, see colored horizontal lines in c). The horizontal dotted line indicates the maximum D’ that can be obtained with random performance. **e**. Self-similarity computed using 1,000 random 2 halves splits of the Go (left) or NoGo (right) trials for the 90°and 15°Go/NoGo angles. Same color code as in (c). **f**. Self-similarity and Go/NoGo similarity for all sessions and Go/NoGo angles. **g**. Activity evoked in the orientation space by the 30°and 60°stimuli of the tuning curve block in trained (top) and naïve (bottom) animals **h**. Average cosine similarity between all pairs of tuning block stimuli in trained (upper triangle) and naive (lower triangle) animals. **i**. Difference between trained and naive mice of the zscored similarity measured for each pair of adjacent stimuli. The shaded area indicates the bootstrapped 95% confidence interval.

### Distortions of orientation representation impact the representational similarity

In a previous study, we had shown that stimuli whose orientations flanked the orientation of the task cues (i.e., ±15°) tended to be represented as the task cues themselves (**Fig. 6g**)(20). Here, we also observed similar distortions of the representation of orientations adjacent to the task cues in trained mice (**Fig. S12-13**) but not in naïve mice (**Fig. S14-15**). To estimate how such distortions affect response similarity, we measured the local cosine similarity i.e., between the responses to adjacent stimuli of the tuning block (e.g., 30°and 60°; **Fig. 6h**). We then compared the local similarity between trained and naïve animals (**Fig. 6i**). We found an increase of similarity for pairs of tuning block stimuli that flanked the orientations of cues used for training, i.e. 30°and 60° (flanking the 45°Go cue, bootstrapped 95% C.I. = [0.49-1.15]) and 120°and 150°(flanking the 135°NoGo cue, bootstrapped 95% C.I. = [0.23-0.74]). This confirmed that stimuli flanking the task cues are represented more similarly, and that the distortions observed in the V1 orientation space have a congruent impact on representational similarity.

### Decoding constrained in orientation space does not outperform the mouse

Our results so far suggested that perceptual discrimination is constrained by the orientation-encoding system and a read-out mechanism that integrates activity from distinct domains of the orientation space. We therefore asked whether a decoder performing a similar integration in the orientation space would outperform the animals in the task, like semantic decoding approaches do (**Fig 1.e-j**). We first trained SNNs to determine the orientations of the 12 tuning block stimuli (**Fig 7a & S16** see methods). After training (**Fig. 7b**), the SNNs were fed with data from the task block. Note that the orientations of the task cues never corresponded to the orientations of the tuning block use as training set, except for the NoGo stimuli presented on D2 (90°) and D6 (60°). The SNNs had 12 output neurons for the 12 orientations of the training set, and the decisions were computed as the circular means of the SNN output vectors. Hence, the decisions could therefore be continuously made in the orientation space. The distributions of the decisions obtained for the Go and NoGo stimuli were used to perform a ROC analysis and provide a D’ (**Fig. S16**). The best possible D’ obtained with this decoding approach matched the animal’s performance (**Fig. 7c**; in descending Go/NoGo angle order, mean±std: 2.92*±*0.77; 2.26*±*0.71; 1.65*±*0.60; 1.43*±*0.57; 1.16*±*0.46; 1.25*±*0.49). Strikingly, the ‘preferred decision weight’ of the V1 neurons (i.e., the circular average of the weights that each neuron had for each of the 12 SNN output, see Methods) ended up corresponding to their preferred orientation computed by using a tuning curve fitting approach (49) (**Fig. S17**). This suggested that the SNNs used neurons in a way that leveraged the same statistical properties of the responses that are captured by traditional tuning curves. These results are compatible with the hypothesis that the read-out of V1 in the context of the task is constrained to the orientation-encoding space.

**Figure 7.**
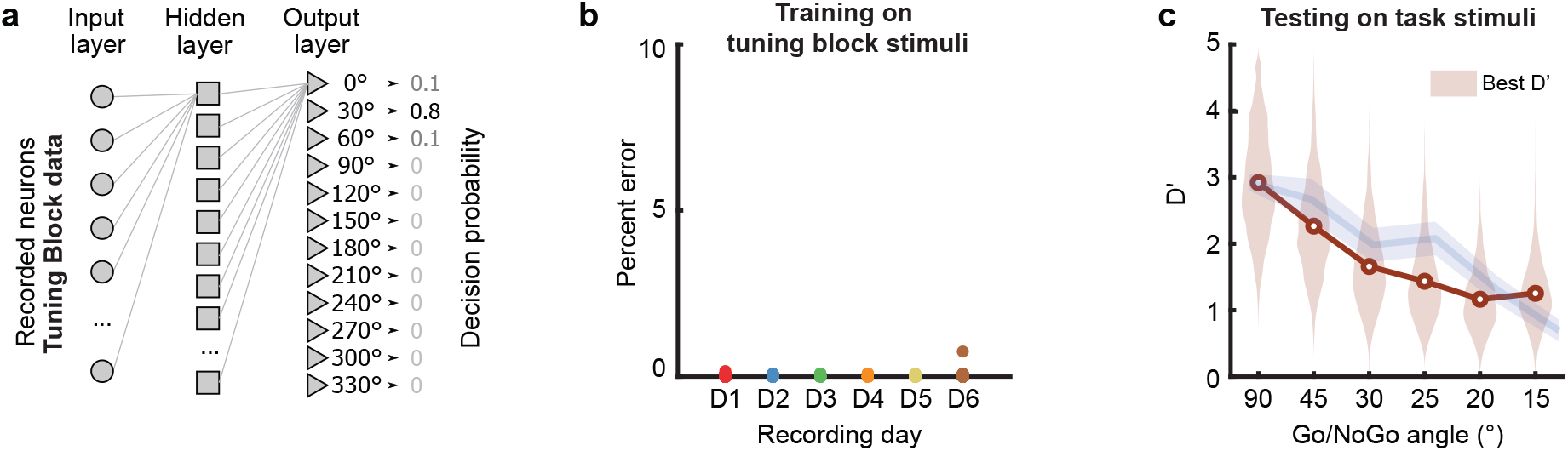
SNNs constrained to the orientation space do not outperform the mouse. **a**. Schematic representation of the shallow neural network trained to classify the orientation of the tuning block stimuli. **b**. Performance of the SNNs at classifying the tuning curve stimuli. **c**. D’ of the same SNNs as in k. when classifying the Go and NoGo stimuli. In brown, the best possible D’ for each SNN computed from a ROC analysis (see Fig. S15). The violin plots show the distribution of values obtained from repeating the classification 1,000 times. In blue is reproduced the animal’s performance as in Fig. 1 for comparison.

## Discussion

The results presented in this study provide several advances in our understanding of the cortical visual processing underpinning perceptual discrimination. By attempting to characterize the relationship between orientation representation and discrimination performance, we unexpectedly revealed that V1 did not faithfully represent orientations during the task. The neuronal activity in the preferred orientation space was not centered on the presented orientation during the task, as it is in naive mice. This result goes against the established idea that V1 faithfully represents orientations in a continuous space. Indeed, around the perceptual threshold, the NoGo cue’s representation was bimodal, with only a relatively weak activation of the neurons tuned for the presented orientation. The two domains of the orientation space co-activated by the NoGo cue were composed of neurons tuned for the orientation of the Go visual stimulus (the Go domain, around 45°) and for orientations at equidistance between the Go and the NoGo stimuli used during training (the NoGo domain, around 90°). We first thought that the presence of those discrete domains of neuronal activity in the orientation space would contribute to decreasing the Go and NoGo representations’ overlap. However, the large activation of the Go domain for the presentation of the NoGo cue ruled out this hypothesis. It instead suggested that the bimodal activations corresponded to a simultaneous probabilistic representation of the two relevant categories of the task, i.e., the Go and NoGo. Indeed, the relative activity at these two domains of the orientation space was found to be a strong predictor of the animal’s choice probabilities. This finding is further supported by the demonstration that changing the relative activation of the Go and NoGo domains, during the presentation of the visual cues in an audiovisual context, results in a predictable improvement of the behavior. Therefore, we have identified for the first time a variable extracted from the V1 output that predicts the performance of the animals. This relative activation of different domains of the orientation space suggests a readout mechanism that integrates the information from ensembles of neurons sharing similar orientation tuning. Such a mechanism can be easily implemented biologically since the connectivity of the early visual processing areas underpins the creation and maintenance of orientation selectivity (50, 12) and its propagation to later stages of processing (51). Instead of weighting the contribution of a few optimally informative neurons, the system likely integrates the output of neurons tuned for relevant orientations, regardless of their reliability and ability to differentiate the stimuli. A selection based on orientation preference (a relatively stable computational property of V1 neurons (52)) rather than abstract information content could protect the readout mechanism from the variability observed in neuronal information content across hours and days (44). Additionally, the orientation-related information was resistant to the variation of the behavioral state of the animals, with the locomotion and pupil size not changing the amount of Go evidence. Importantly, we found that the discrete activation of the Go and NoGo domains was not reflecting the behavioral choices of the animals. Indeed, the trial-to-trial Go and NoGo representation variability did not correlate with the animal’s choice. The modulation observed when comparing correct and incorrect trials was orthogonal to the Go evidence. This suggests that V1 provides probabilistic estimates of the categoric membership of the stimuli, and that behavioral choice variability is created in subsequent processing stages (44, 4). This finding of a probabilistic encoding of the visual cue is in line with inferential theories of sensory processing, such as the hierarchical Bayesian inference hypothesis of visual processing (53). Modeling and specific inquiry of the integrating process at work here are needed to further understand the mechanisms underlying the relationship between V1 output and discrimination performance we uncovered. Finally, we found that the Go and NoGo domains are shaped by a modulation of the neuronal excitability in the orientation space that we did not observe in naïve mice passively viewing the stimuli. This effect was symmetrical in that it was present around both 45°and 135°in the preferred orientation space, implying it is not dependent upon the difference of rewards and behaviors associated with the Go and NoGo trials. We had previously shown that training could induce such modulations through a surround suppression around the task stimuli (20). We also show, using a qualitative model, that the bimodal response in the orientation space could be explained by the multiplicative interaction between an input centered on the visual stimulus orientation and the modulation of the neuronal excitability in the orientation space. Additionally, the persistent location of the NoGo domain around 90°could be a consequence of the spatial extent of the surround suppression that arises around the Go (45°) and NoGo (135°) cues during training. The cellular mechanisms underpinning the representation changes found in trained mice remain to be determined. Surround suppression is an ubiquitous mechanism associated with shaping sensory representations (54). However, how such a surround suppression can be established in a feature-encoding space such as orientation is pending further investigation. It is likely that the preferred-orientation-specific suppression observed in trained mice relies on interneuron-mediated inhibition. During learning, interneurons are modulated and facilitate cortical plasticity (55, 56, 57, 58, 59, 60), and they receive several top-down signals (61, 62, 63, 59). Yet, dopaminergic and cholinergic neuromodulation could also be involved as they are also linked to perceptual learning and cortical plasticity (64, 65, 66, 67, 68) via specific modulation of V1 microcircuit’s actors. Importantly, the suppression of flanking neurons was not completely context-dependent, as it was evidenced during the tuning block when mice were passively viewing the stimuli. Moreover, despite the 135° NoGo stimuli losing relevance after the first recording day, the suppression of neurons representing 135°-flanking orientations persists through the six consecutive days of recording. The surround suppression is therefore likely to involve local plasticity (16), but could be a transient state of V1 that reverts to a non-distorted orientation space when the task loses relevance (20). Future research should investigate the interplay between top-down modulation, local synaptic plasticity, GABAergic inhibition, and neuromodulation in order to reveal the circuit underpinnings of the orientation space distortion generated during perceptual training.

## Methods

All the procedures described below have been approved by the Institutional Animal Care and Use Committee of Rutgers University, in agreement with the Guide for the Care and Use of Laboratory Animals.

### Surgery

This study used adult (3-6 months old) male and female mice in three datasets. The main dataset consisted of 10 C57BL6 mice (Jackson Laboratories #000664). The second dataset with unimodal visual and audiovisual conditions consisted of 10 Gad2-IRES-Cre (Jackson Laboratories #019022) × Ai9 (Jackson Laboratories #007909) mice. The third dataset with naïve mice passively viewing stimuli consisted of 7 Ai163 x Camk2a-Cre (Jackson stock #005359) mice, expressing GCaMP6s in excitatory cells. Ten minutes after systemic injection of an analgesic (carprofen, 5 mg per kg of body weight), mice were anesthetized with isoflurane (5% induction, 1.2% maintenance) and placed in a stereotaxic frame. Body temperature was kept at 37°C using a feedback-controlled heating pad. Pressure points and incision sites were injected with lidocaine (2%). Eyes were protected from desiccation with artificial tear ointment (Dechra). Next, the skin covering the skull was incised, and a custom-made lightweight metal head-bar was glued to the skull using Vetbond (3M). In addition, a large recording chamber capable of retaining the water necessary for using a water-immersion objective was built using dental cement (Ortho-Jet, Lang). A circular craniotomy (diameter = 3 mm) was performed above V1. For the main and audiovisual datasets, the AAV vector AAV1.eSyn.GCaMP6f.WPRE.SV40 (UPenn Vector Core) carrying the gene of the fluorescent calcium sensor GCaMP6f was injected in three sites separated by 500 μm around the center of V1 (stereotaxic coordinates: -4.0 mm AP, +2.2 mm ML from bregma) using a MicroSyringe Pump Controller Micro 4 (World Precision Instruments, WPI) at a rate of 30 nL/min. Injections started at a depth of 550 μm below the pial surface and the tip of the pipette was raised in steps of 100 μm during the injection, up to a depth of 200 μm below the dura surface. The total volume injected across all depths was 0.7 μL. After removal of the injection pipette, a 3-mm-diameter coverslip was placed over the dura, such that the coverslip fits entirely in the craniotomy and was flush with the skull surface. The coverslip was kept in place using Vetbond and dental cement. During the 5 days following surgery, mice were provided amoxicillin administered in drinking water (0.25 mg/mL). Mice were left to recover from the surgery for at least 3 weeks to obtain a satisfactory gene expression.

### Behavioral training

Mice were water-deprived up to 85% of their body weight and acclimated to head fixation on a spherical treadmill in a custom-built, soundproof training rig. The rig was equipped with a monitor (Dell) and a water dispenser with a custom infrared beam lickometer. Data acquisition boards (National Instruments and Arduino) were used to actuate the solenoids for water delivery and vacuum reward retrieval, as well as recording the animal licking activity. The monitor and data acquisition boards were connected to a computer that ran a custom-made training program scripted in MATLAB. Once animals reached their target weight and were acclimated to the training setup, they were trained to perform a Go/NoGo orientation discrimination task (Fig. 1A). Go/NoGo tasks are classically used to investigate the mechanisms underpinning perception and perceptual learning in rodents. They rely on a common perceptual discrimination behavior that consists in discriminating a desirable stimulus from non-desirable ones (69, 24). The direct association between stimulus, response and reward limits the impact of cognitive and/or executive errors inherent to more complex tasks, allowing a more straightforward readout of the animals’ orientation discrimination abilities. In our task, drifting sinewave gratings oriented 45° below the horizontal were paired with a water reward, and the animal was expected to lick (Go trial). Drifting gratings orthogonal to the Go signal (orientation 135°) signaled the absence of reward, and the animal was expected to withhold licking during those trials (No-Go trial). The Go and NoGo cues were presented full screen to cover the full visual field of the recorded hemisphere, allowing to maximize the number of neurons excited by the stimuli and make sure that the receptive fields of the recorded neurons would remain on the stimulus despite changes in gaze direction. Those drifting gratings had the same temporal frequency (2 Hz), spatial frequency (0.04 cycle per degree), duration (3 seconds) and contrast (75%). When the stimulus instructed the animal to lick, the water delivery was triggered by the mouse licking during the third second of the stimulus presentation. No water was dispensed in the no-lick condition, or if the mouse failed to trigger water delivery in the lick condition. If the animal responded correctly (Hit or Correct Rejection), the intertrial interval was 3s. If the animal responded incorrectly (Miss or False Alarm), the intertrial interval was increased to 6s as a negative reinforcement (Fig. 1A). Performance was measured using the D’ statistic (D’=norminv(Hit rate) norminv(False alarm rate), norminv = inverse of the normal cumulative distribution function). Animals were considered experts if their performance during training sessions was greater than 1.7.

#### Best random D’

Our training strategy was that mice first acquired a strongly reliable licking behavior for the Go, then learned to withhold from licking in NoGo trials. Hence, the improvement of their performance during training was primarily driven by a decrease in the False Alarm rate. To account for the constant bias of mice toward licking, we computed the best random D’ value using a probability of licking in response to the Go cue that corresponded to the average Hit rate over the six recording sessions. The best random D’ was defined as the point where the mouse would differentiate the NoGo cue from the Go cue in half of the trials, i.e., a FA rate of 50%. For the first dataset, we therefore considered a binomial distribution of p = 0.93 and n = 128 (average number of trials) for the Go behavior, and of p = 0.5 and n = 128 for the NoGo behavior. We used the 77.5 and 22.5 percentiles to simulate the best hit rate and FA rate that could be obtained (p=0.05) given these underlying probabilities. This led to a value of D’ = 1.68. For the second dataset, where animals licked less for the Go when the Go/NoGo angle decreased, and the number of trials was lower, we used a binomial distribution of p = 0.77 and n = 60 for the Go and p = 0.5 and n = 60 for the NoGo, yielding a value of D’ = 1.0. These values correspond to the D’ values above which we can conclude that mice do not behave at random for the NoGo trials, not to the perceptual threshold.

### Imaging sessions

#### First dataset (unimodal visual trials only)

Calcium imaging in V1 was performed over 6 consecutive days. During those sessions, the orientation of the Go signal remained at 45°, while the orientation of the NoGo signal was set to 135° for the first session, 90° for the second, 75°for the third, 70°for the fourth, 65°for the fifth, and 60°for the sixth. Hence the angle between the Go and the NoGo cues was progressively decreased: Day 1: 90°, Day 2: 45°, Day 3: 30°, Day 4: 25°, Day 5: 20° and Day 6: 15° (Fig. 1A). Our decision to record with different NoGo values for six consecutive days was motivated by the following reasons. [1] We needed the mouse behavior to be modulated by the perceptual difficulty not the task difficulty. Our goal was therefore to make sure that mice continuously understood the task they were trained for. By decreasing gradually the angle between the Go and NoGo cue we made sure that the mouse maintained their understanding of the task rules. The mice result in being more exposed to the task for the late sessions (e.g., when the Go/NoGo angle is 15°) than for the early sessions (e.g., when the Go/NoGo angle is 45°). However, this is unlikely to affect the conclusion we can draw from our results. [2] The Go signal was also kept constant across sessions to simplify the task rules for the mouse and allow for a better estimate of the mouse orientation discrimination threshold (decreasing the chance that the animal does not perform because it does not understand the task rules). Moreover, this decreased the number of free parameters in our experiments, as the NoGo orientation is the only parameter that changes across the completion of the task. [3] Finally, as V1 activity is very sparse and highly variable from trial to trial, we also wanted to work with a large number of trials for each Go/NoGo angle. Mice typically perform a maximum of 200 trials per training session. However, to make sure that mice were optimally motivated to perform the task while we were recording, we kept the number of trials to 120 to avoid misses and correct rejection trials due to a lack of motivation to perform the task.

#### Second dataset (unimodal visual and audiovisual trials)

In this dataset (collected by a different experimenter), calcium imaging in V1 was performed during 5 consecutive daily sessions using the same visual stimuli as described above for the first dataset starting from a Go/NoGo angle of 45°. The only difference was that the sessions were divided into 4 blocks of 30 trials alternating unimodal visual and audiovisual blocks (see below; Fig. 3A) At the beginning of the recording session, the first block was randomly selected between unimodal visual and audiovisual.

#### Third dataset (naïve animals)

In this dataset, mice were recorded for six consecutive days and passively exposed to the same sequence of visual stimuli as the mice from the first dataset (Fig. S4). Out of 7, 5 were exposed to the stimuli in the same order (decreasing angle between the two gratings) and 2 were exposed in the reverse order (increasing the angle between the two gratings).

#### Recording sessions

Recordings were made of three types of blocks: unimodal visual, audiovisual (only for the second dataset), and tuning blocks. For all those blocks, the gratings had the same temporal frequency (2 Hz) and spatial frequency (0.04 cycle per degree). The duration was 3 seconds for the unimodal visual and audiovisual blocks and 1.5s in the tuning blocks. The contrast of the drifting gratings was 25% during testing (compared to 75% during training) to enhance the impact of sounds on representation of the visual stimuli orientation in V1 (sound modulation in V1 was shown to follow the principle of inverse effectiveness (43, 45). Auditory stimuli during the audiovisual blocks consisted of the presentation of one of two sinewave pure tones (10 kHz and 5 kHz; 78 dB; duration: 3 seconds). Sounds were produced by a speaker located immediately below the center of the screen. Visual and auditory stimuli were generated in Matlab (MathWorks) using the Psychtoolbox (70). The background noise (room heat and air conditioning airflow, laser cooling fans, and spherical treadmill airflow) was 69 dBA. Each audiovisual trial resulted from the random combination of one of the two pure tones with one of the two drifting gratings of the session. For each imaging session, we assessed the orientation tuning of the imaged neurons in an orientation tuning block that consisted in the presentation of a series of drifting sinewave gratings (12 orientations evenly spaced by 30 degrees and randomly permuted) passively viewed by the mice. The tuning block was usually located at the end of the imaging session. However, placing the tuning block at the beginning of the recording session did not change our estimation of the neurons preferred orientation (Fig. S1). The spatiotemporal parameters of tuning block stimuli were identical to those of the behavioral block except for their duration and contrast (temporal frequency = 2 Hz, spatial frequency = 0.04 cycle per degree, contrast = 100%; duration: 1.5 seconds; intertrial interval: 3 seconds). As scanning was not synchronized with the stimulus presentation, a photodiode located at the top left corner of the screen was used to detect the exact timing of the visual stimuli onset and offset. The photodiode signal was acquired along with the following signals: 1) a signal provided by the two-photon microscope, which indicated the onset of each frame, and 2) two analog signals encoding the orientation of the drifting grating and the frequency of the auditory stimulus. These signals were digitized (NiDAQ, National Instruments) and recorded with the software WinEDR (John Dempster, University of Strathclyde). Ball motion was tracked by an IR camera taking pictures of the ball at 30 Hz. Eye motion was monitored at 15 Hz using a second IR camera imaging the reflection of the eye on an infrared dichroic mirror. Functional imaging was performed at 15 frames per second using a resonant scanning two-photon microscope (Neurolabware) powered by a Ti-Sapphire Ultra2 laser (Coherent) set at 910 nm. The scanning mirrors of the Neurolabware microscope are in a hermetically sealed chamber, bringing the scanning hum below the room ambient noise (< 59 dBA). The laser beam was focused 200 microns below the cortical surface using a 16×, 0.8 NA Nikon water-immersion objective. The objective was tilted 30 degrees such that the objective lens was parallel to the dural surface. Laser power was kept below 70 mW. Frames (512×796 pixels) were acquired using the software Scanbox developed by Neurolabware.

### Data analysis

#### Imaging data pre-processing

Calcium imaging frames were realigned offline to remove movement artifacts using the Scanbox algorithm (Neurolabware). For the first dataset (see Table S1), a region of interest (ROI) was determined for each neuron using an automatic segmentation and pre-processing routine (Suite2P (71)). The second dataset (see Table S1) was segmented by hand. For every frame, the fluorescence level was averaged across the pixels of the ROI. Potential contamination of the soma fluorescence by the local neuropil was removed by subtracting the mean fluorescence of a 2-5-micron ring surrounding the neuron’s ROI, excluding the soma of neighboring neurons, and then adding the median value across time of the subtracted background. We then computed the fractional fluorescence from the background subtracted fluorescence data. The fractional fluorescence (dF/F = (F – F0) / F0), was calculated with F0 defined as the median of the raw fluorescence measured during every inter-trial interval, then was deconvolved into Action Potential related Events (APrEs) using a benchmarked deconvolution algorithm (69, 24). Spike inference has several advantages over directly using the calcium signal. The calcium signal intrinsically provides an indirect estimate of the neuron’s activity. Action potentials induce a sharp increase in intracellular calcium concentration followed by a slower decay. This decay deteriorates the temporal resolution, and consecutive action potentials add up nonlinearly. Additionally, the noise due to artifacts or baseline drift is sometimes the same order of magnitude as the spiking activity. This leads to an imprecise estimate of the neuronal activity. Spike inference reduces the impact of these limitations, allowing to improve the temporal resolution, to account for the nonlinearity of spike signal addition, and to reduce the impact of noise. Only neurons whose correlation between the inferred fractional fluorescence (convolved back from the inferred APrEs) and the measured fractional fluorescence was greater than 0.8 were used for the analysis. This inclusion criterion removed neurons for which the calcium signal was too poor to allow the algorithm to accurately detect APrEs. Additional inclusion criteria were used for the automatically segmented dataset (‘iscell’ variable from suite2p > 0.8, minimum area > 15 *μm*^2^, roundness (% of overlap with a circle centered on the ROI) > 20, and response probability for the preferred orientation of the Tuning Block >0.1). Trials were isolated and stored on a SQL database. Some imaging sessions in the second dataset yielded data that could not be used because of poor viral expression or unusable cranial windows, but the behavioral data for these animals was kept (see Table S1).

#### Tuning curves

The orientation tuning curve of each neuron was fitted to the APrE responses recorded during the tuning curve blocks using a Markov chain Monte Carlo sampling-based Bayesian method (20, 49, 43) using a Poisson noise model. This algorithm fitted 4 different models to the observed responses (“circular gaussian 360°” for orientation but not direction selective tuning curves, “circular gaussian 180°” for strictly direction selective tuning curves, “direction selective circular gaussian” for direction and orientation tuned neurons, and “constant” for untuned neurons). The fit that explained best the data was then selected. If the constant model was the best fit, the neuron was considered not tuned. The preferred orientation was defined as the peak of the fitted orientation tuning curve. When a neuron was not direction selective (i.e., responding equally to the same oriented stimulus moving in opposite directions), the preferred orientation was defined as the orientation included in the range [0 – 180°].

#### Similarity

For each recording session, the mean activity across time of the n simultaneously recorded neurons was represented by an n-dimensional vector. The cosine similarity was computed as the cosine of the angle between the activity vectors evoked by the Go and NoGo trials. The same procedure was then repeated using all neurons from all sessions to obtain a grand average. The point having the largest deviation from the baseline value during the first second of the stimulus presentation was used to capture the relationship between similarity and performance. Baseline similarity was defined as the mean similarity measured during the 1 second period immediately preceding the stimulus presentation.

#### Representation of the neuronal activity in the orientation space

To allow a full coverage of the orientation space, neurons were pooled across similar recording sessions (for a same Go/NoGo angle and a same dataset). The preferred orientation of every neuron was assessed (using the fits of the APrE responses recorded during the tuning block, see above). A subset of ROIs of the first dataset did not show significant orientation or direction tuning when we did not apply the inclusion criteria (18.18%, n = 12602/69317). However, when the inclusion criteria were applied (i.e., ‘iscell’ parameter from Suite2P, roundness, minimum area and response probability), all of the neurons showed a reliably estimated orientation or direction tuning. In the second dataset, a large subset of the segmented ‘neurons’ did not show significant orientation or direction tuning (46.89%, n = 1667/3555), and were excluded from further analysis. For each stimulus of each group (e.g., 45° presented to trained mice), we generated an orientation space activity map, sorting neurons according to their preferred orientation. Their activity was then averaged across trials, baseline-subtracted, and averaged across neurons sharing the same preferred orientation. To access the variability across trials of the pooled pseudo-population, we generated 1,000 resampled trials using the following procedure. For each resampled trial, 250 neurons (15% of the neurons of the database) were randomly selected, and for each neuron, one trial from the test block during which the stimulus of interest was presented (e.g., 45°) was selected at random. The activity of the neuron during the selected trial was included in an orientation*time matrix in which rows corresponded to the neurons’ preferred orientations (bin: 1°). The resulting trial matrix was baseline-subtracted and smoothed across the orientation space by a Gaussian weighted mean (std= 6°).

#### Locomotion and pupil size

Locomotion (ball motion) and pupil size were recorded and analyzed for the dataset including the audiovisual version of the task, since it was necessary to show that sound-induced representation changes were not attributable to change in locomotion or pupil size. The raw traces were lowpass filtered (0.25 Hz) and then either normalized on their intra-session standard deviation (locomotion) or z scored (pupil). For pupil size, a small number of trials were discarded because either the mouse was blinking, or the position of the eye was not allowing us to reliably assess the size of the pupil. We compared the amounts of both behavioral variables within sessions with a permutation test, and between session averages with a paired Wilcoxon test. To evaluate the impact of the quantity of locomotion and pupil size on orientation representations, we sorted all trials in quartiles according to the amount of locomotion/pupil size and compared the Go evidence values, since they depend on the distribution of activity in the orientation space.

#### Mean tuning curves comparison

To compare the tuning curves of neurons in specific bins of orientation preference, the tuning curve fit of all considered neurons was circularly shifted so that the peak would be centered on 0°. To visualize the data underlying the averaged fits, the APrE responses were shifted by the same number of degrees. For example, the data from a neuron with a preferred orientation of 45°was shifted by -45°such that its response to 0°would show at -45°on the plot, its response to 30°would show at -15°, etc. The median of each individual neuron’s responses was then computed, and for each degree of the orientation space the medians from all neurons falling in that degree (that were equally shifted) were averaged.

#### Tuning curve amplitude across orientation space

The amplitudes of the tuning curve fit all neurons from sessions of the first dataset having the same Go/NoGo angle were sorted according to their preferred orientation (bin width = 7.5°) and averaged for each bin. The same operation was performed including all the neurons from all the sessions of the first dataset, and the result was fitted using a smoothing spline (MATLAB fit function, smoothing parameter = 0.01). Using different smoothing methods (rolling average, gaussian weighted rolling average) or bin sizes did not change the results (not shown). The mean location of the peak and troughs and their confidence intervals were obtained with bootstrapping the tuning curve amplitude data 1000 times, fitting the smoothing spline, and locating the local maxima and minima.

#### Bimodality at the trial level

We represented the different hypotheses by simulating random Gaussian distributions in two dimensions (MATLAB function normrnd) for the “exclusive” and “constant” hypotheses, and a random Pearson distribution (MATLAB function pearsrnd) for the “correlated” hypothesis. To assess whether the coactivation of the Go and NoGo domains observed in the average of the pseudo-population was present within an individual session and at the trial level, we analyzed the population activity of single trials in the Go/NoGo domains space. For every trial, the APrEs evoked during the two first seconds of stimulus presentation by all the neurons with preferred orientation ranging between 1° and 60° (Go domain) or 61° and 120° (NoGo domain) were summed and divided by the number of neurons considered. For each trial, we then plotted the activity measured in the Go domain versus the activity measured in the NoGo domain. An ellipse was fitted on that cloud so it would encompass 95% of the trials in that space. The orientation of the ellipse being aligned with the orientation of largest variance of the point cloud, it was used to compare the variance measured in every session to the expected variance orientation that would be observed if a Go/NoGo coactivation would not happen at the trial level. Additionally, the optimal number of clusters to explain the data was evaluated with the Calinski-Harabasz criterion (MATLAB evalclusters function).

#### V1 representation vs. decision at the trial level

To test whether the clouds of single trial activity in the Go/NoGo domains space were different when trials were split according to the behavioral decision of the animal, we used a two-dimension extension of the Kolmorogov-Smirnov test (72). The direction of the cloud displacement between successful and unsuccessful trials was assessed by drawing a line between the centroid of the two clouds.

#### Shallow neural networks (SNN)

We used SNNs to classify stimuli-evoked responses for three different types of analyses. 1.) To determine if V1 neuronal populations contain the information necessary to outperform the mice at the task. For each recording session (10 mice * 6 Go/NoGo angles) we randomly selected 250 neurons, or as many as available, and randomly ordered trials with a rng seed so within-trial correlations are maintained. The SNNs were trained and tested using the sum of APrEs evoked during the stimulus presentation. Seventy percent of the trials were used for training, 10% for validation and the remaining 20% were used for testing. SNNs were made of an input layer that corresponded to the recorded neurons, a hidden layer (n=150 neurons) and an output layer (n=2). We used the MATLAB function patternnet, with a scaled conjugate gradient backpropagation or “trainscg” algorithm. The number of training epochs was set to 300 and the training goal to 1e-5. The regularization parameter was set to 0.2 to limit overfitting. This procedure was repeated 100 times for each recording session. For every one of these repetitions, we also created 10 other SNNs trained using randomly label-shuffled trials as a control. We also performed a similar analysis using pseudo-populations. To do so, we randomly picked 1000 sets of 250 neurons from the whole dataset and created pseudo trials by bootstrapping 100 trials for every neuron. The rest of the parameters were the same as described above. 2.) To evaluate the decoding performance of a SNN when operating in the feature space. We trained SNNs with pseudo-populations (see above) using the activity recorded during the tuning block. The output layer was made of 12 neurons, one for each orientation in the tuning block set (from 0° to 330°, evenly spaced). Neurons were selected from the whole dataset and the procedure was repeated 1000 times. Once trained, the SNNs were tested using a 100 bootstrapped trials array for the same neurons, but this time using trials recorded during the task block (Go and NoGo). The output vector of the SNN (12 values, one for each output neuron) provided a probability distribution. The orientation of the presented stimulus estimated by the SNN was defined as the circular-mean of that output vector in the 0-180°range. For each SNN, we tested the SNN with 100 Go and NoGo trials, and computed ROC curves by sliding a decision criterion across the 100 orientations estimated by the SNN. The D’ were computed by finding the best possible performance, i.e. the ROC curve point furthest from the bisector. 3.) To extract the orientation information present in the responses of every neuron as an alternative to our classical tuning curves pipeline. SNNs were trained in the feature space using the tuning block data as described above (SNN application #2). For each trained SNN, we computed the influence of every input neuron on the SNN output decisions (*W*_*input→output*_) using the connection weight method (73). This resulted in a 12-value vector (one weight for each output). The preferred weight orientation was computed as the circular average of the connection weight vector wrapped to 180°. Wrapping was necessary to avoid orientation-aligned high weights cancelling each other and non-orientation-aligned high weights averaging to an orientation that doesn’t correspond to the neurons’ information content. Neurons could be selected multiple times when generating the 1,000 SNNs. When it was the case, we computed the circular average of their preferred weight orientation provided by the SNNs that used them. The preferred weight orientation of each V1 neuron was then compared to the preferred orientation computed by fitting the orientation tuning curve (Fig. S17).

#### Statistics

To determine if the means of two distributions (amplitude of tuning curves, behavioral responses, and D’) were significantly different, we compared the observed difference with the distribution of 10,000 differences obtained when the labels were shuffled. The two-tailed confidence interval of the null hypothesis at the alpha level 0.05 was defined as the 2.5 and 97.5 percentile of the distribution obtained from the permutations. The difference between the observed means was considered significant if located outside the confidence interval of the null distribution. Because large number of observations can lead to conclude on significant differences of even irrelevantly small magnitude, we complimented relevant tests with an effect size measure, Hedges’g, using a MATLAB toolbox (74). Traditionally, values under 0.2 correspond to the absence of effect, values between 0.2 and 0.4 are considered to reflect a small effect, values between 0.4 and 0.6 correspond to a medium effect, and values larger than correspond to a large effect size. In some cases, when comparing smaller samples (skewness of Go representation, Go Evidence as a function of locomotion and pupil size quartiles), we used a paired Wilcoxon test with an alpha level of 0.05. The relationship between cosine similarity and D’ was fitted using the MATLAB Central File Exchange function sigm_fit (R P (2023). (https://www.mathworks.com/matlabcentral/fileexchange/42641-sigm_fit). The pseudotrials’ peak distributions were fitted with a mixture of von Mises distributions using mvmdist (https://github.com/chrschy/mvmdist), a Matlab package for probabilistic modeling of circular data. This allowed to localize the largest components in the orientation space. The correlations between Go evidence and choice probability were computed using Pearson correlations (MATLAB function corrcoeff). The isotropy of the decision-induced displacement of the activity in the Go evidence space (Fig. 4) was assessed using Watson’s U test. The effect of the Go/NoGo angle on representation self-similarity was assessed using one-way ANOVAs. Responsive neurons (Fig. S5) were identified as in (20) using a binomial test that compared the probability of having at least one APrE during the stimulus presentation to the probability of having at least one APrE during the prestimulus time.

## Notes

### Competing Interest Statement

The authors have declared no competing interest.

### Summary of Updates

The manuscript has been extensively revised. It includes additional experiments (naive mice) and novel complementary analyses (shallow neural networks and cosine similarity).

