## Supplementary Material for "Discretized representations in V1 predict suboptimal orientation discrimination"

**The PDF file includes:**

Materials and Methods
Supplementary Text
Figs. S1 to S17
Tables S1

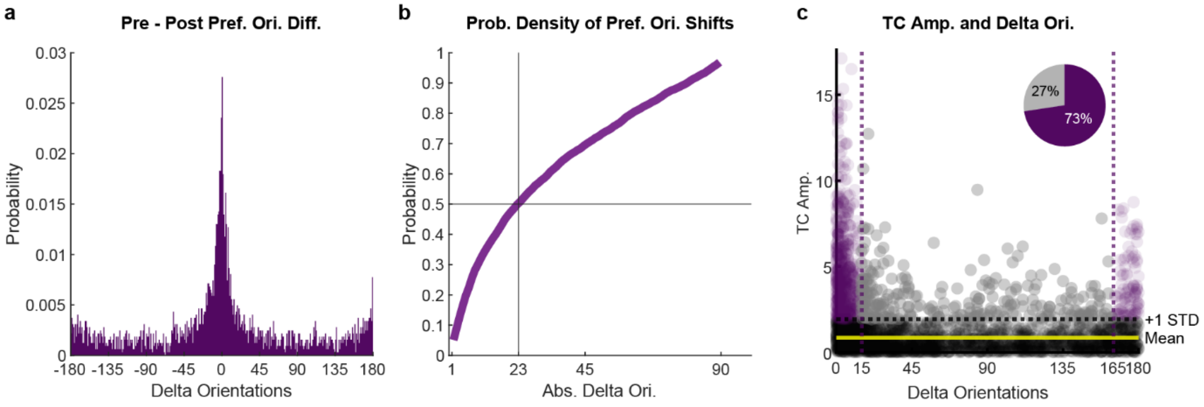


**Fig. S1. Stability of V1 neurons' preferred orientations during the recording sessions. a.** Distribution probability of the shift in orientation preference (delta orientation) estimated during the tuning block preceding (pre) or following (post) the behavioral task (n = 3 mice). The distribution is centered on 0, indicating a stability of the preferred orientation during the recording session. **b.** Probability distribution of the absolute value of the orientation shift (Abs. Delta Ori.). 50% of the neurons having a preferred orientation had a preferred orientation shift of less than 23° (median shift). **c.** Amplitude of the response of the neurons to their preferred orientation (TC amp.: tuning curve amplitude) as a function of their shift in orientation preference during the recording session. We computed the mean and standard deviation (SD) of the tuning curve amplitude of neurons with the largest shifts (between 15° and 165°) and defined a threshold as the mean + 1SD of their preferred orientation response. 73% of the neurons with a tuning curve amplitude greater than this threshold had a preferred orientation shift of less than 15° (purple neurons). Therefore, most of the most responsive neurons and thus the most likely to participate in the representation of the task cues had a preferred orientation shift of less than 15°.

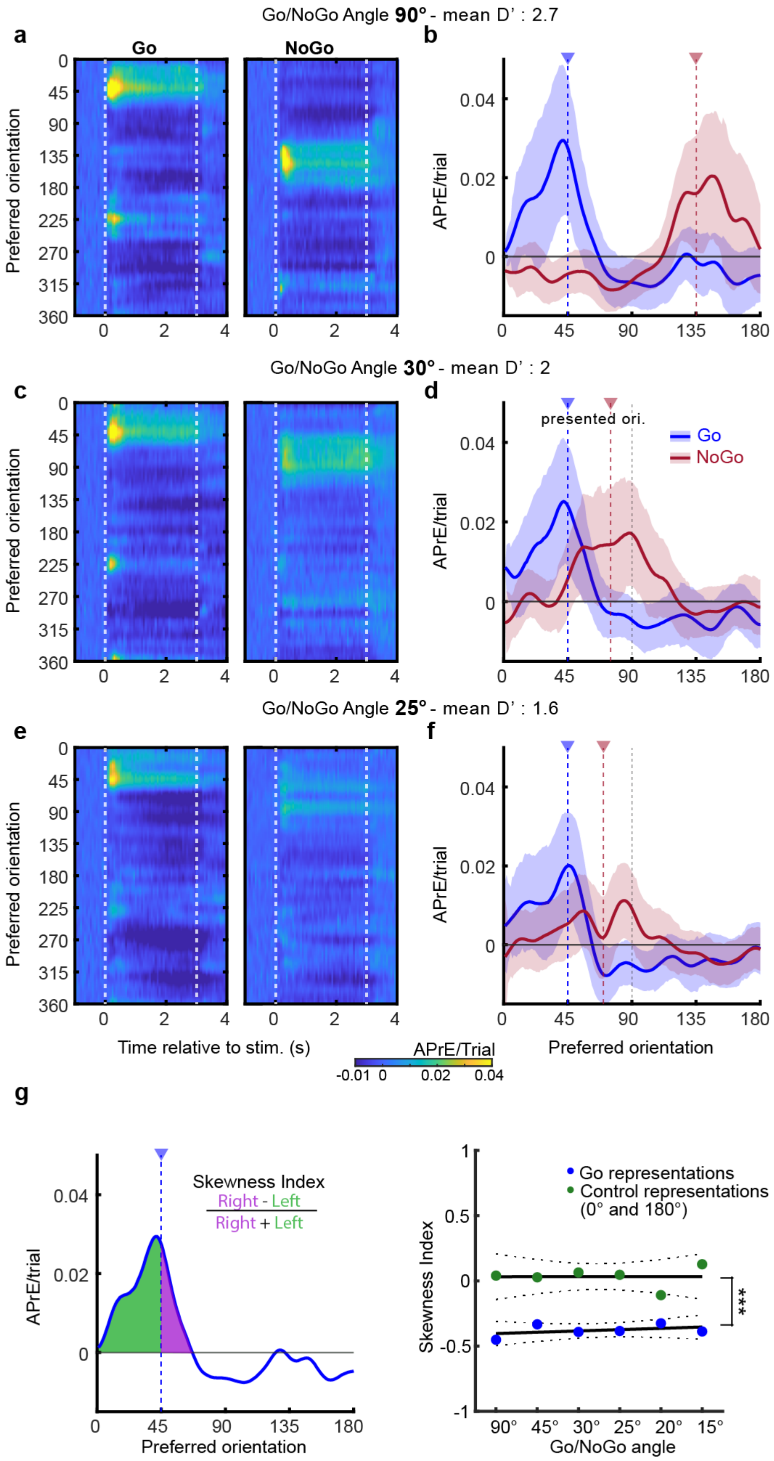


**Fig. S2. Representations of Go and NoGo stimulus in the V1 orientation space for the other Go/NoGo angles (first dataset, unimodal visual). a.** Neuronal activity evoked by the Go (left panel) and NoGo stimuli (right panel) as a function of V1 neurons' preferred orientations when the Go/NoGo angle is 90°. Neuronal activity color scale is located at the bottom of the (e) panel. The mean behavioral performance (D') across mice is indicated on top. **b.** Mean activity evoked in the orientation space by the Go (45°, blue) and NoGo (135°, red) stimuli during the two first seconds of the stimulus presentation. Vertical colored lines indicate the orientation of the presented stimulus. Shaded areas indicate the s.e.m. **c.** Same representation as in (a) when the angle between the Go/NoGo angle is 30°. **d.** Same representation as in (b) when the NoGo orientation is 75°. **e.** Same representation as in (a) when the Go/NoGo angle is 25°. **f.** Same representation as in (b) when the NoGo orientation is 70°. **g.** Left: Schematic of the skewness index computation. Right: Skewness of Go representations vs. control orientations representation (0 and 180°). The skewness values for the Go representations were negative and significantly different from 0, indicating a higher density of activity on the side of the orientation space away from the NoGo orientation (mean+- S.E.M. = -0.38 +- 0.02 Wilcoxon test, n=6 p=0.028). This skewness of the stimulus representation in the orientation space was not found for the control orientations 0° and 180° (skewness index = 0.03 +- 0.03, p = 0.25).

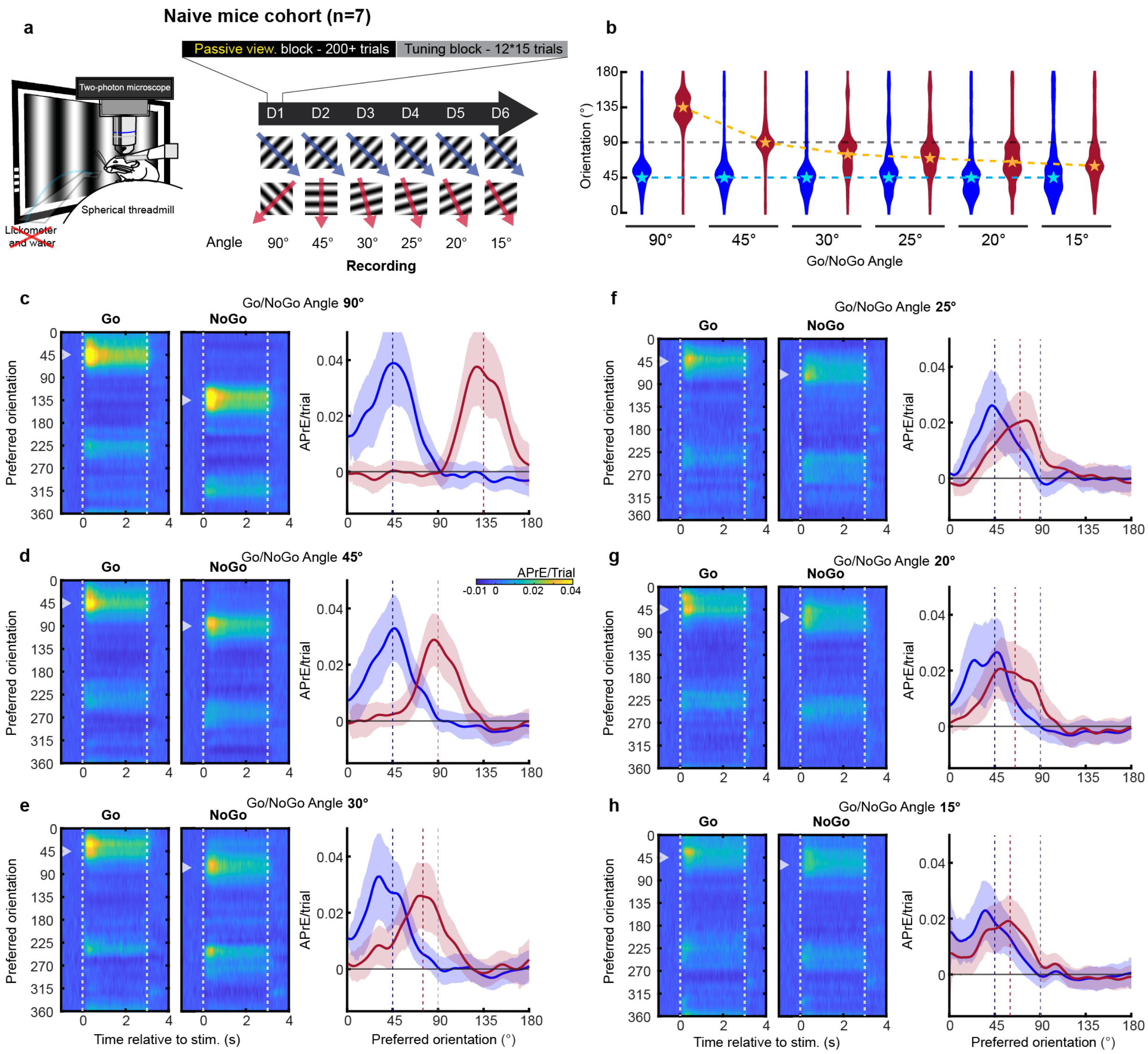


**Fig. S3: Orientation representation in naïve animals passively viewing the task stimuli. a.** Schematic representation of the experimental setup and the organization of the recording sessions (top right) including the Go and NoGo stimuli used for the different recording days (bottom right). **b.** Distribution of the location of the peak of activity in the orientation space evoked by the Go (blue) and NoGo (red) pseudo-trials (n=1000, see Methods). Blue and gold stars respectively indicate the orientation of the Go and NoGo stimuli presented during the task. **c-h.** Action Potential related Events (APrEs) generated by the presentation of the Go (45°, left panel) and NoGo stimuli (135° to 60°; Go/NoGo cue angles: 90° to 15°, right panel) as a function of the V1 neurons' preferred orientation (average across all neurons having the same preferred orientation).

**Fig. S4: Bimodal representations of the NoGo cues during single trials. a.** Schematic description of the tested hypotheses (see Methods). (i) The bimodal representations of the NoGo cue found for Go/NoGo angles between 20° and 30° when averaging across trials (black dotted lines) could result from the activation of the neurons tuned for the Go domain (gold) and neurons tuned for the NoGo domain (magenta) in different trials ("exclusive hypothesis", top left panel). If it were the case, plotting for each trial the activity evoked in the NoGo domain against the activity evoked in the Go domain would lead to a plot with two distinct clusters (bottom left panel). (ii) The bimodal representations could also result from the simultaneous activation of the Go and NoGo domain in each single trial ("consistent hypothesis": top right panel). In that case, the Go and NoGo domain would be similarly activated across trials leading to a single cluster (bottom right panel). (iii) Finally, this bimodal representation could result from an intermediate hypothesis, where for each trial the activation of Go domain is anti-correlated to the activation of NoGo domain ("correlated hypothesis": top center panel). In that case, we would find a single cluster elongated along an axis perpendicular to the line of equity (axis angle: 45°; see inset of bottom center panel). **b.** Domains' activation of all the trials of a session of the first dataset during which the Go/NoGo angle was 20°. Note that NoGo trials are represented by a single cluster. The the NoGo cues activate similarly Go and NoGo domains (cluster on the line of equity). **c.** Domains' activation of all the trials of another session during which the Go/NoGo angle was 25°. Note the similarity of those results with those presented in (b). **d.** Response profiles across the 2 first second of the NoGo stimulus presentation (left panel) for all the sessions of the *first dataset* when the Go/NoGo angle was 20°. Those profiles lead to single clusters (central panel) which maximum variance axes ranges between 0° and -90° (right panel). This result disproves the exclusive and anti-correlated hypotheses that would result in a variance oriented around +45°. **e.** Same representation as in (d) for the all the sessions of the *second dataset* when the Go/NoGo angle was 25°. **f.** Distribution of the maximal variance orientation of the domain activation evoked by the presentation of the NoGo cue for all the Go/NoGo angles supporting the "consistent hypothesis". Kruskal Wallis: $H(5)=25.76$; $p = 0.0001$. **g.** In most sessions, neuronal activity evoked by the Go (top panels) and NoGo (bottom panels) single trials generate a single cluster in the domain activation space (optimal number of cluster evaluated with the Calinski-Harabasz criterion) in the Go/NoGo domains activation space, disproving the "exclusive hypothesis". Altogether, those analyses demonstrate that the NoGo and Go domains are co-activated by the NoGo stimuli at the single trial level when the Go/NoGo angle is near the limit of perceptual discrimination.

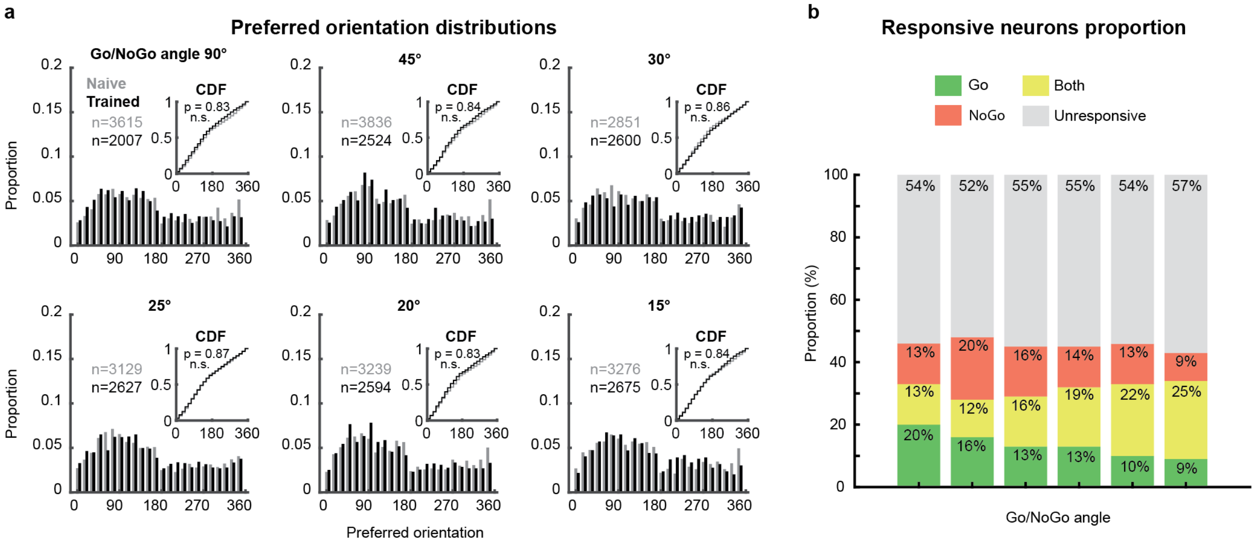


**Fig. S5: Distribution of the preferred orientations and proportion of responsive neurons in L2/3 V1 neurons. a.** Distribution of the preferred orientations of all recorded neurons in Trained (black) and Naïve animals (grey). The insets show the cumulative density functions for both groups. The p values were obtained by an Anderson-Darling test comparing the trained and naive distributions. **b.** Proportion of neurons responding to the Go, NoGo or both stimuli. The proportion of unresponsive neurons remains similar across days.

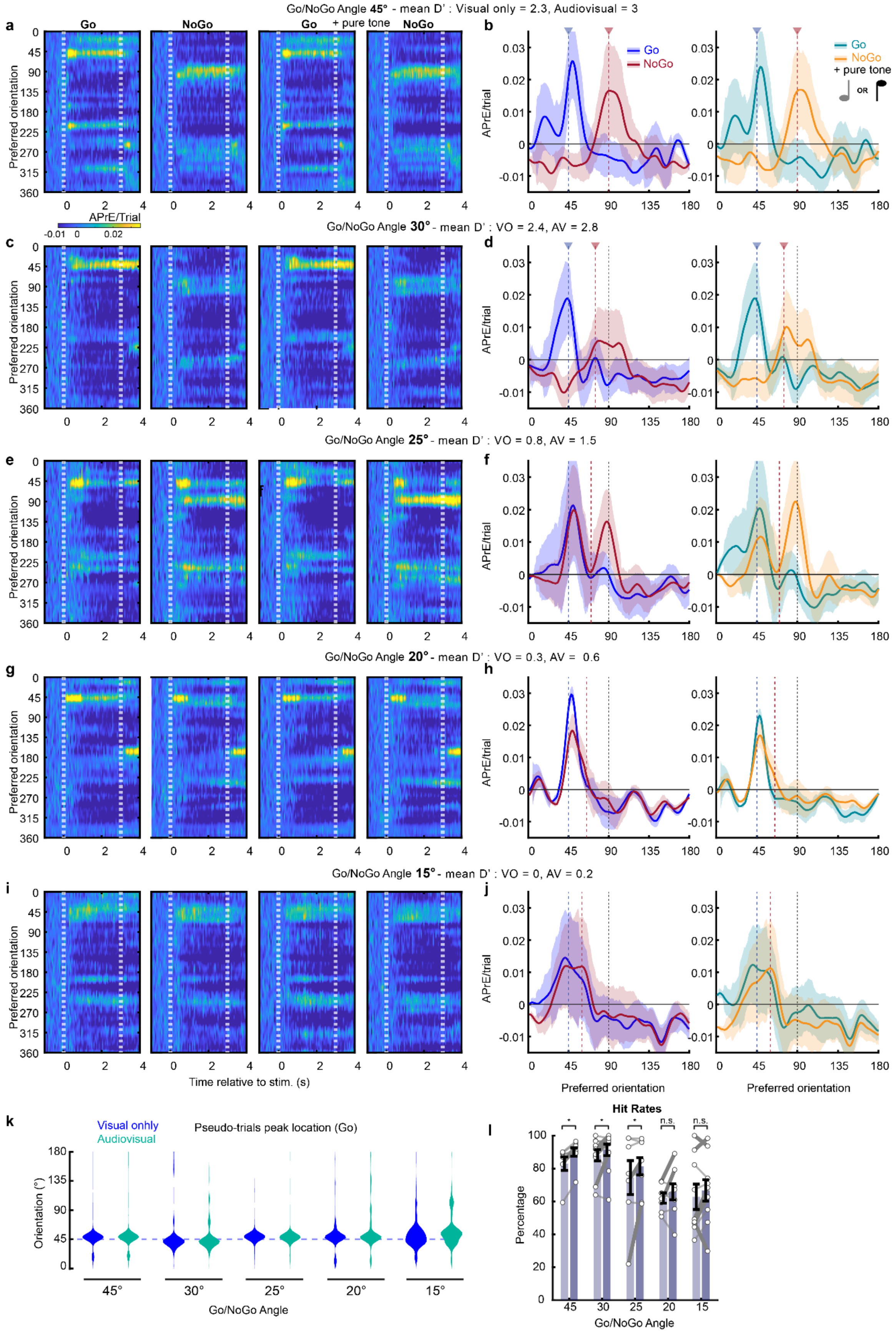


**Fig. S6. Representations of the Go and NoGo stimuli in the orientation space as a function**

**of the Go/NoGo angle for the second dataset (with unimodal visual and audiovisual blocks).** **a.** Neuronal activity evoked by the Go (first panel on the left) and NoGo stimuli (second panel on the left) as a function of V1 neurons' preferred orientations when the Go/NoGo angle is 45° in the unimodal visual blocks. Right panels: same representation for the Go (third panel) and NoGo stimuli (fourth panel) presented in the audiovisual blocks. Neuronal activity color scale is located at the bottom of the panel. The mean behavioral performance (D') across mice is indicated on top. **b.** Left panel: mean activity evoked in the orientation space by the Go (45°, blue) and NoGo (90°, red) stimuli during the two first seconds of the stimulus presentation in the unimodal visual blocks. Right panel: mean activity evoked in the orientation space by the Go (45°, cyan) and NoGo (90°, orange) stimuli during the two first seconds of the stimulus presentation in the audiovisual blocks. Vertical colored lines indicate the orientation of the presented stimuli. Shaded areas indicate the s.e.m. **c.** Same representation as in (a) when the angle between the Go/NoGo angle is 30°. **d.** Same representation as in (b) when the NoGo orientation is 75°. **e.** Same representation as in (a) when the Go/NoGo angle is 25°. **f.** Same representation as in (b) when the NoGo orientation is 70°. **g.** Same representation as in (a) when the Go/NoGo angle is 20°. **h.** Same representation as in (b) when the NoGo orientation is 65°. **i.** Same representation as in (a) when the Go/NoGo angle is 15°. **j.** Same representation as in (b) when the NoGo orientation is 60°. **k.** Violin plot of the distribution of the location in the orientation space of the peak of activity for unimodal Go (blue) and audiovisual Go (cyan) pseudo-trials. Dotted line indicates the orientation of the presented Go stimulus. **l.** Hit rates during the unimodal visual (light purple) and audiovisual (dark purple) blocks for the different Go/NoGo angles. Black lines indicate mean ± s.e.m. across sessions. Statistical tests: ns: $p>0.05$; *: $p<0.05$; ***: $p<0.001$. White dots indicate individual mice (thick lines indicate significant difference; $p<0.05$).

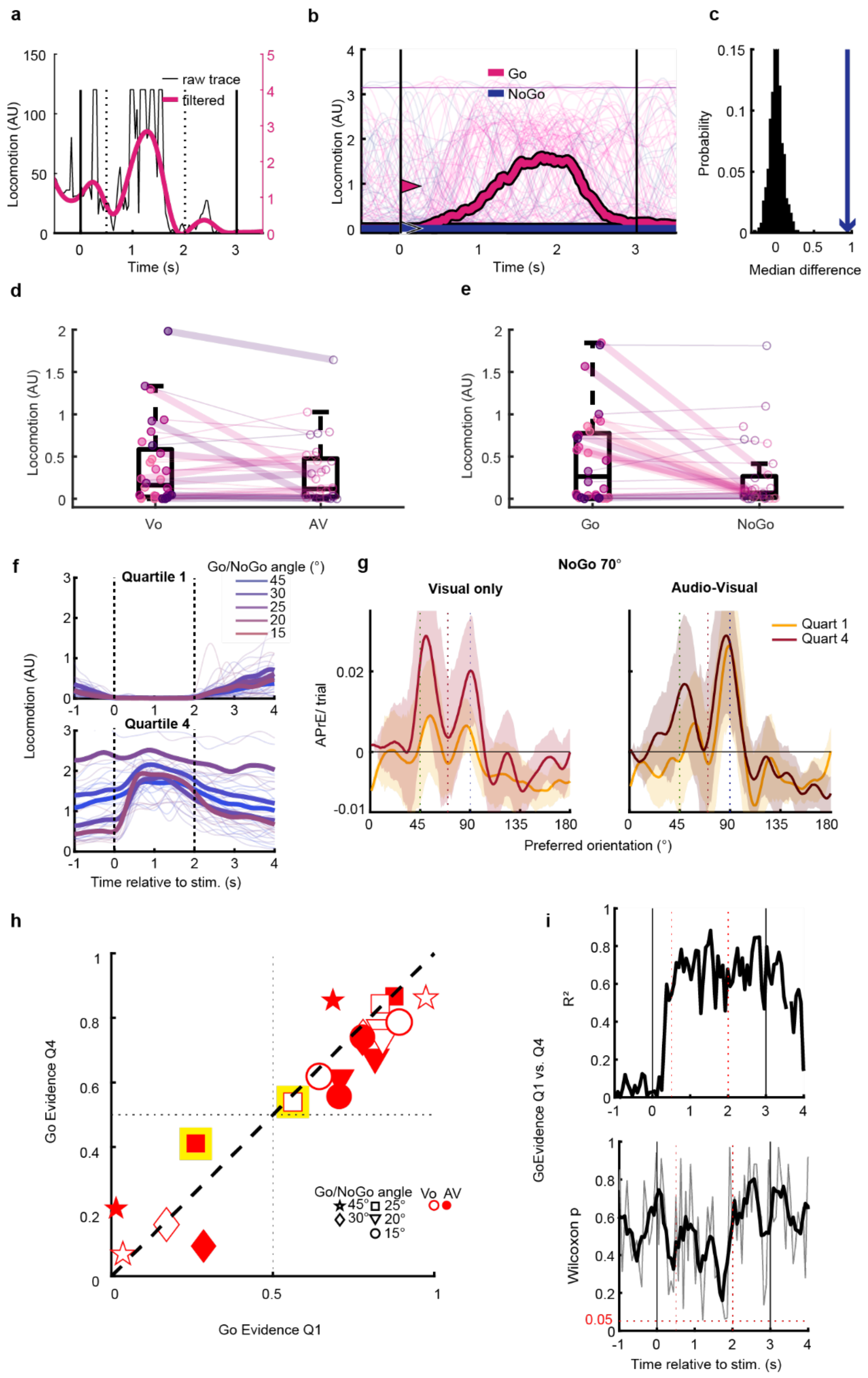
a
b
c
Locomotion (AU)
raw trace
filtered
Time (s)
Go
NoGo
Probability
Median difference
d
e
Vo
AV
Go
NoGo
f
Quartile 1
Quartile 4
Go/NoGo angle (°)
Time relative to stim. (s)
g
NoGo 70°
Visual only
Audio-Visual
Quart 1
Quart 4
APrE/ trial
Preferred orientation (°)
h
Go Evidence Q4
Go Evidence Q1
Go/NoGo angle
Vo
AV
i
GoEvidence Q1 vs. Q4
R²
Wilcoxon p
0.05
Time relative to stim. (s)

**Fig. S7. Locomotion does not explain orientation representation distortions nor the relationship between V1 representation and behavior. a.** Black trace: measure of the locomotion speed during a single trial (in arbitrary units). Purple trace: low pass filtered trace. **b.** Locomotion measured for all the trials of a single session separating the Go (pink) and NoGo (blue) trials. Thin lines indicate single trials. Thick lines the averages across trials. Arrowheads indicate the average amount of locomotion during the analyzed epoch (i.e., between 0.5 and 2s, indicated by dashed lines). **c.** Permutation test comparing the observed difference in average locomotion (blue) against the distribution of differences found when shuffling the Go and NoGo labels from b. (black) **d.** Locomotion measured for all sessions in the Visual Only (Vo) and audio-visual (AV) conditions. While 15.6% of sessions had a significant difference (permutation test $p<0.05$) in locomotion between Vo and AV conditions, the difference was not found at the cohort level (paired Wilcoxon test, $p=0.45$). **e.** Locomotion measured for all the sessions during the presentation of the Go and NoGo stimuli. While 31.3% of sessions had a significant difference (thick lines, permutation test $p<0.05$) in average locomotion, the cohort did not have a significantly different amount of locomotion between the two conditions (paired Wilcoxon test, $p=0.064$). This indicated that some but not all mice tended to run in Go trials, a behavior that we observed when they were licking to get the reward. **f.** Average locomotion for trials belonging to the first (top) and fourth (bottom) quartiles of locomotive activity. Quartiles were computed according to the locomotion average between 0 and 2s post stimulus. Thin lines are individual sessions, thick lines indicate the average of all sessions for the different Go/NoGo angles. **g.** Average activity profiles evoked by the 70° NoGo in the orientation space for trials in the first (yellow) and fourth (red) quartiles of locomotive activity, in the Visual Only (left) and Audio-visual (right) conditions. Bimodal activity profiles were found in all the cases. Locomotion seemed to be associated with a change in the amplitude but not the orientation-space distribution of the evoked activity. **h.** Comparison of the “Go Evidence” measured in the first and last locomotion quartiles showing that the Go evidence is similar in the presence or absence of locomotive activity. The yellow highlights show the values measured in the activity profiles shown in g. **i.** The $R^2$ values of the Go Evidence correlations between quartiles 1 and 4 remain above 0.5 (average = 0.68) between 0.5 and 2s post-stimulus, and the paired Wilcoxon test p values remain above 0.05 (average = 0.44) during that same epoch, demonstrating that the locomotion does not significantly change the Go Evidence.

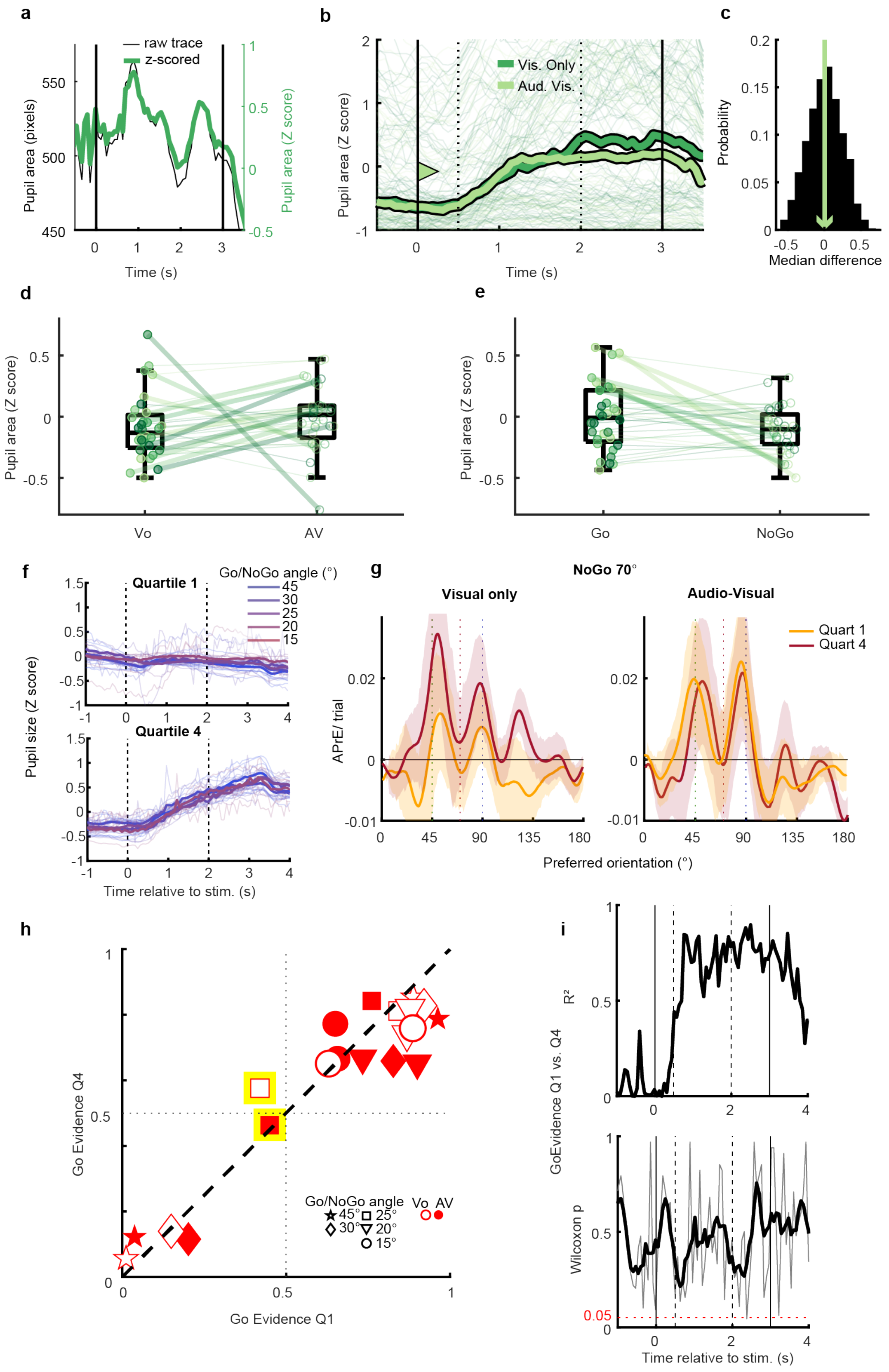
a
raw trace
z-scored
Pupil area (pixels)
Pupil area (Z score)
Time (s)
b
Vis. Only
Aud. Vis.
Pupil area (Z score)
Time (s)
c
Probability
Median difference
d
Pupil area (Z score)
Vo
AV
e
Pupil area (Z score)
Go
NoGo
f
Quartile 1
Go/NoGo angle (°)
45
30
25
20
15
Quartile 4
Pupil size (Z score)
Time relative to stim. (s)
g
NoGo 70°
Visual only
Audio-Visual
Quart 1
Quart 4
APrE/ trial
Preferred orientation (°)
h
Go Evidence Q4
Go Evidence Q1
Go/NoGo angle
45°
25°
30°
20°
15°
Vo
AV
i
R²
GoEvidence Q1 vs. Q4
Wilcoxon p
0.05
Time relative to stim. (s)

**Fig. S8. Arousal does not explain orientation representation distortions nor the relationship between V1 representation and behavior. a.** Variation of the pupil size during a single trial, in pixels (black) and Z-scored (green). **b.** Average pupil area for an example session for the Visual only (dark green) and audio-visual (light green) conditions. Thin lines indicate single trials. Arrowheads indicate average pupil size during the analyzed epoch (i.e., between 0.5 and 2s, indicated by dashed lines). **c.** Permutation test comparing the observed difference in average (green) against the distribution of differences when shuffling the visual only and audiovisual labels. (black) **d.** Comparison of the pupil area between visual only and audiovisual blocks for all sessions. While only 12.5% of sessions presented a significant difference (permutation test $p<0.05$) in pupil area between those conditions, the cohort had higher pupil area in the audiovisual condition (paired Wilcoxon test, $p=0.02$). **e.** Comparison of the pupil area measured during the Go and NoGo trials for all sessions. While 25% of sessions presented a significant difference (thick lines, permutation test $p<0.05$) in pupil area between the two cues, the pupil area was similar between the two conditions at the cohort level (paired Wilcoxon test, $p=0.16$). The similarity of the pupil size for the two visual stimuli suggests that the orientation representation distortions observed for the NoGo stimuli do not depend on pupil size. **f.** Average pupil area measured for trials belonging to the first (top) and fourth (bottom) quartiles of pupil area. Quartiles were computed according to the average pupil area measured between 0 and 2s of the stimulus presentation. Thin lines indicate individual sessions, thick lines indicate the average of all sessions for the different Go/NoGo angles. **g.** Average activity profiles evoked in the orientation space by the 70° NoGo cue for trials in the first (yellow) and fourth (red) quartiles of pupil area, for the Visual Only (left) and Audio-visual (right) conditions. Bimodal activity profiles were found in all cases. Arousal seems therefore to be associated with a change in the amplitude but not the distribution of the evoked activity in the orientation space. **h.** Comparison of the "Go Evidence" measured during the two extreme quartiles since that variable depends on the distribution of activity in the orientation space. The average Go Evidence throughout 0.5 to 2s post-stimulus lies around the unity line. The yellow highlight shows the values derived from the profiles in g. **i.** The $R^2$ values of the Go Evidence correlations between quartiles 1 and 4 remain above 0.5 (average = 0.68) between 0.5 and 2s post-stimulus, and the paired Wilcoxon test p values remain above 0.05 during that same epoch (average = 0.38), showing that arousal does not significantly change the Go Evidence.

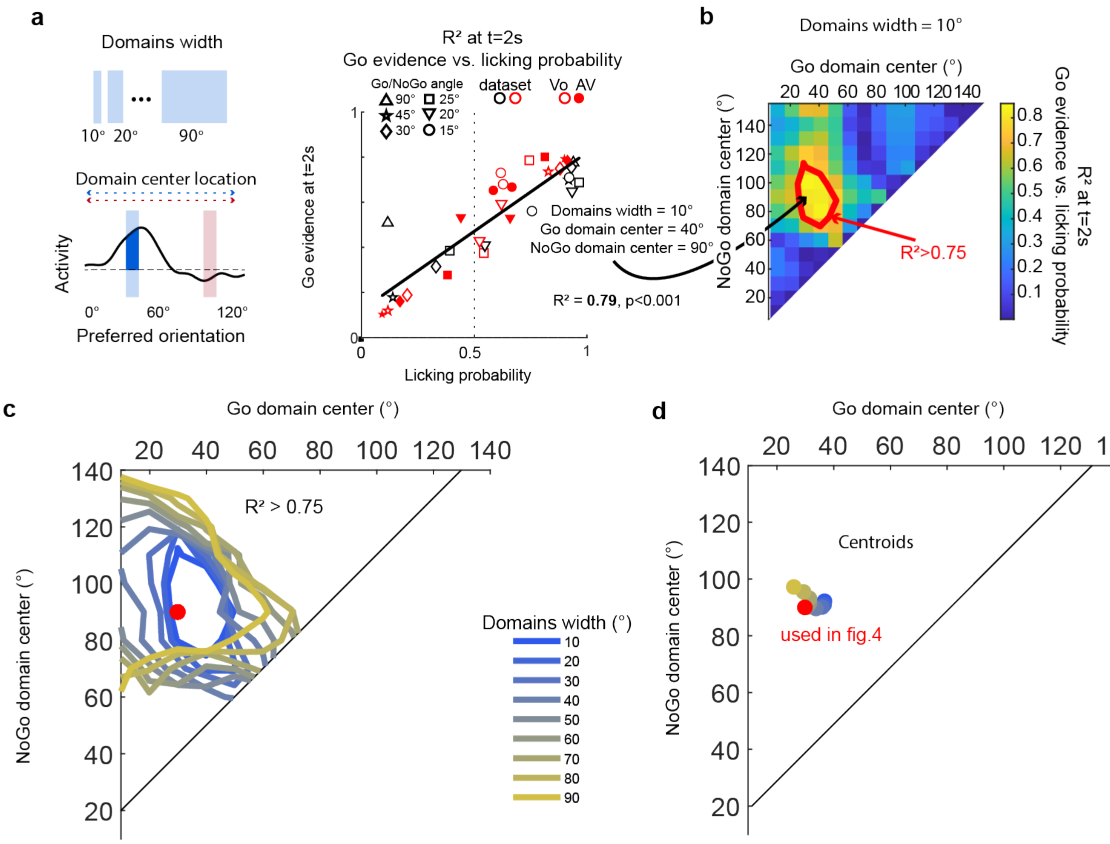


**Fig. S9. Systematic evaluation of different locations and widths for the Go and NoGo domains. a**. Schematic presentation of the procedure. The width of the Go and NoGo domains were systematically increased in 10° increments between 10 and 90°, and the domains' center was placed between 10 and 150° in 10° increments. The data from the evoked profiles of activity for all combinations of domains' widths and locations was used to correlate the Go Evidence (domain 1 activity / (domain 1 + domain 2 activity)) with the licking probability. This allowed to relate the domains' parameters with the strength of the correlation between Go Evidence and behavior. **b.** $R^2$ of the correlations between Go Evidence and licking probability for all couples of domains' locations and a width of 10°. The red polygon indicates the region in the location space for which the R>0.9. In that case, domains centered around 40 and 90° are the best at predicting the licking probability from the V1 activity. **c.** Contour of the location-space regions for which the correlation between Go evidence and licking probability has a $R^2$ value>0.9, for each tested width. **d.** Centroids of those regions. All domain widths can yield similar high correlation strengths, as long as they encompass exclusively the 40° and 90° regions of the orientation space. Therefore, the domains' parameters used in figure 4 are optimal to capture the relationship existing between the activity profiles in the orientation space and the behavior.

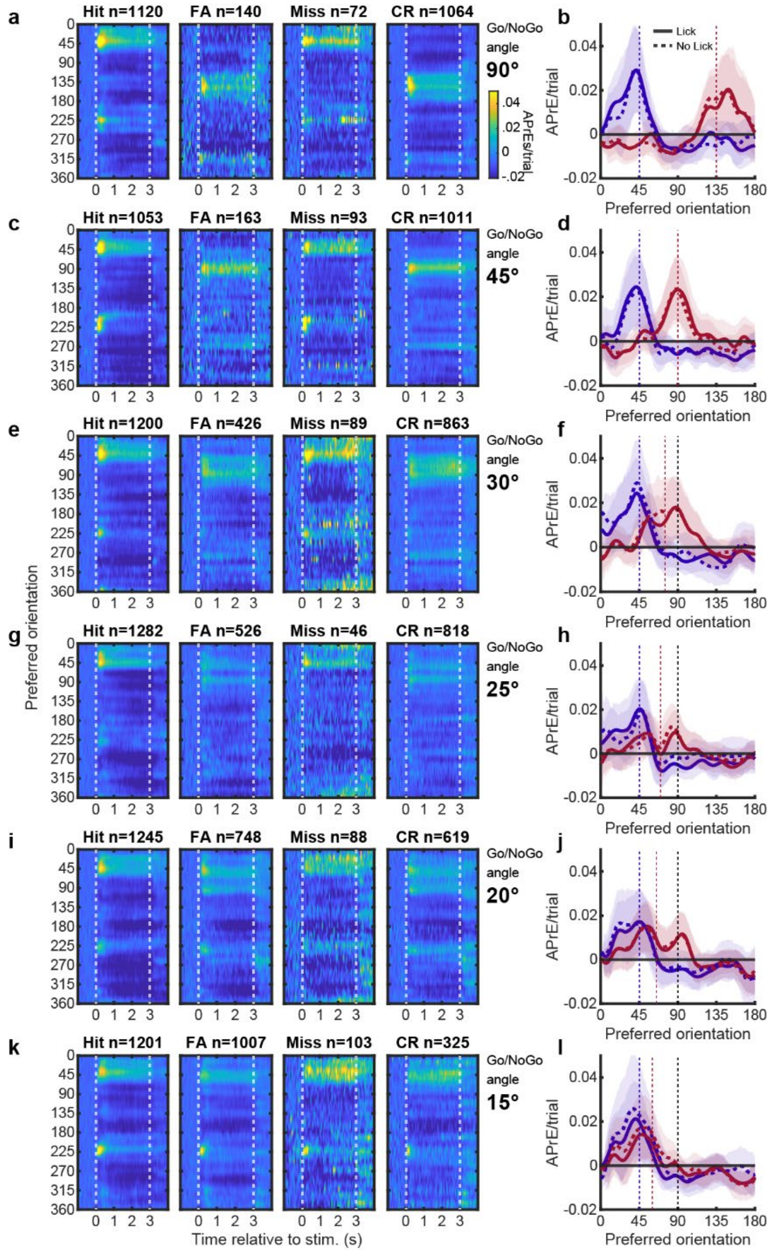


**Figure S10: Comparison of Go and NoGo cue representation in trials split by behavioral decision. a.** Neuronal activity evoked by the Go and NoGo stimuli as a function of V1 neurons' preferred orientations when the Go/NoGo angle was 90°. The top text indicates whether the trials used for each plot were Hits, False Alarms (FA), Misses or Correct rejection (CR), as well as the number of trials available for each type. **b.** Mean activity evoked in the orientation space by the Go (45°, blue) and NoGo (135°, red) stimuli during the two first seconds of the stimulus. Plain lines indicate licking decisions, i.e. Hits and FAs. Dashed lines indicate the absence of licking during the rewarded window, i.e. Misses and CR. **c-d.** Same as (a) and (b) but for the 45° Go/No angle. **e-f.** Same as (a) and (b) but for the 30° Go/No angle. **g-h.** Same as (a) and (b) but for the 25° Go/No angle. **i-j.** Same as (a) and (b) but for the 20° Go/No angle. **k-l.** Same as (a) and (b) but for the 15° Go/No angle.

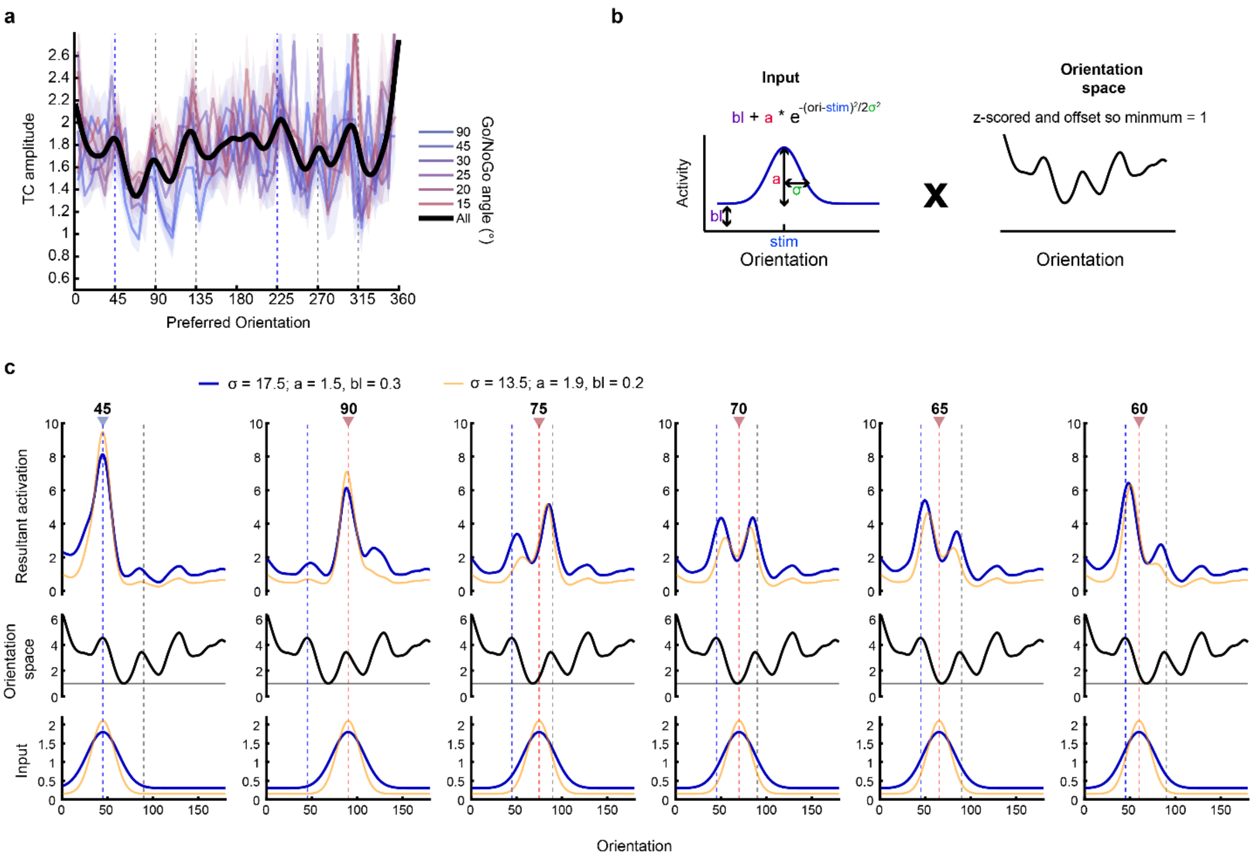


**Fig. S11: Putative mechanism underpinning the bimodal representation of the NoGo cues in the orientation space. a.** Modulation of the neuronal excitability in the orientation space (0-360°) for the different Go-NoGo angles. Black trace: averaged modulation across sessions. A modulation of the neuronal excitability in the orientation space is also found for neurons preferring gratings drifting in the opposite direction. **b.** Testing the effect of a multiplicative interaction between a unimodal activation in the orientation space by the visual stimulus (modelling a 'pure' bottom input to layer 2/3; left panel) and the modulation of the neuronal excitability in the orientation space found experimentally in layer 2/3 (right panel). **c.** Effect (top panels) of the multiplicative interaction between a unimodal visual activation of the orientation space (bottom panels) and the modulation of the neuronal excitability in the orientation space (middle panel) for each Go/NoGo angle (columns). Vertical lines indicate the orientation of the presented stimuli (blue: Go orientation, purple: NoGo orientations). Blue traces: inputs and outputs corresponding to the presentation of the stimulus in the unimodal visual context. Yellow traces: inputs and outputs corresponding to the presentation of the stimulus in the audiovisual context. The effect of sound on the inputs (decrease of baseline, increase of peak and decrease of width) was qualitatively modeled after observations reported in Ibrahim et al., 2016. Those results qualitatively reproduce the activity profiles obtained experimentally.

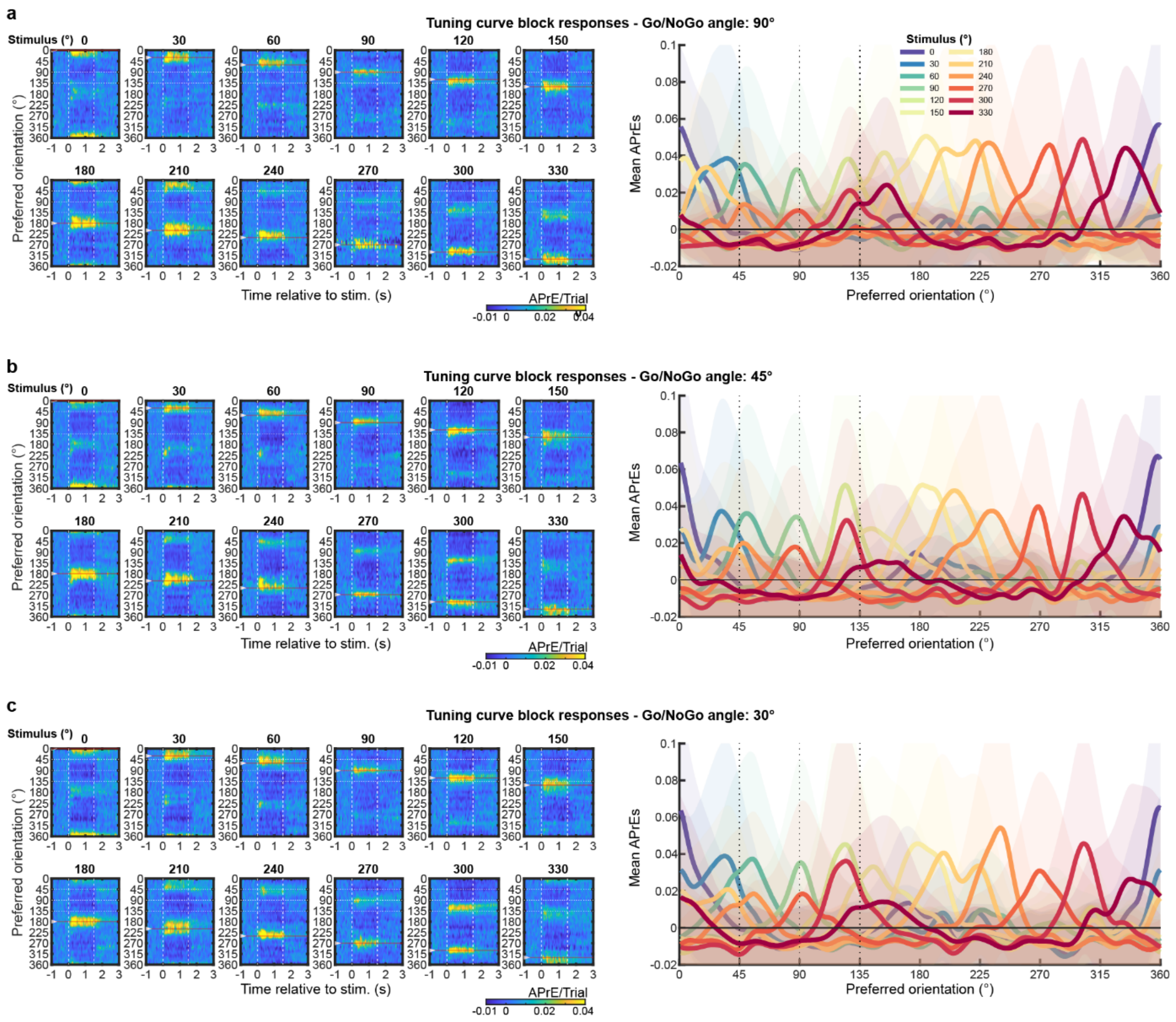


**Fig. S12: Representations in V1 of tuning block stimuli in trained animals between recording days 1-3. a.** *Left*: Neuronal activity evoked by the presentation of all 12 tuning block stimuli as a function of the neurons' preferred orientations. *Right*: Average activity profile in the preferred orientation space during the two first seconds of stimulus presentation for all stimuli of the tuning block following the recording session D1. Mean ± s.e.m. **b.** Same representation as in a. for the recording session D2. **c.** Same representation as in a. for the recording session D3.

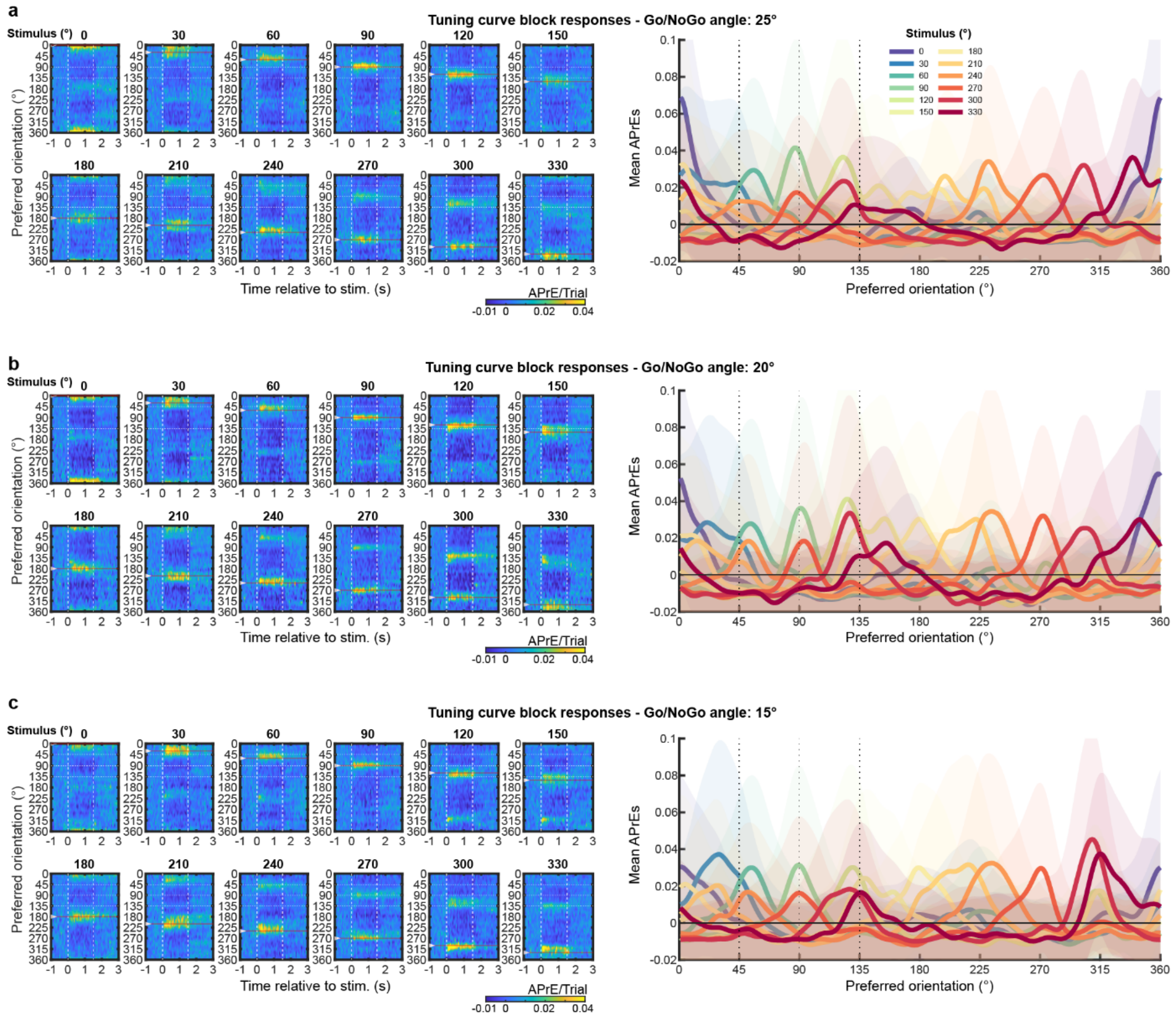


**Fig. S13: Representations in V1 of tuning block stimuli in trained animals between recording days 4-6. a.** *Left*: Neuronal activity evoked by the presentation of all 12 tuning block stimuli as a function of the neurons' preferred orientations. *Right*: Average activity profile in the preferred orientation space during the two first seconds of stimulus presentation for all stimuli of the tuning block following the recording session D4. Mean ± s.e.m. **b.** Same representation as in a. for the recording session D5. **c.** Same representation as in a. for the recording session D6.

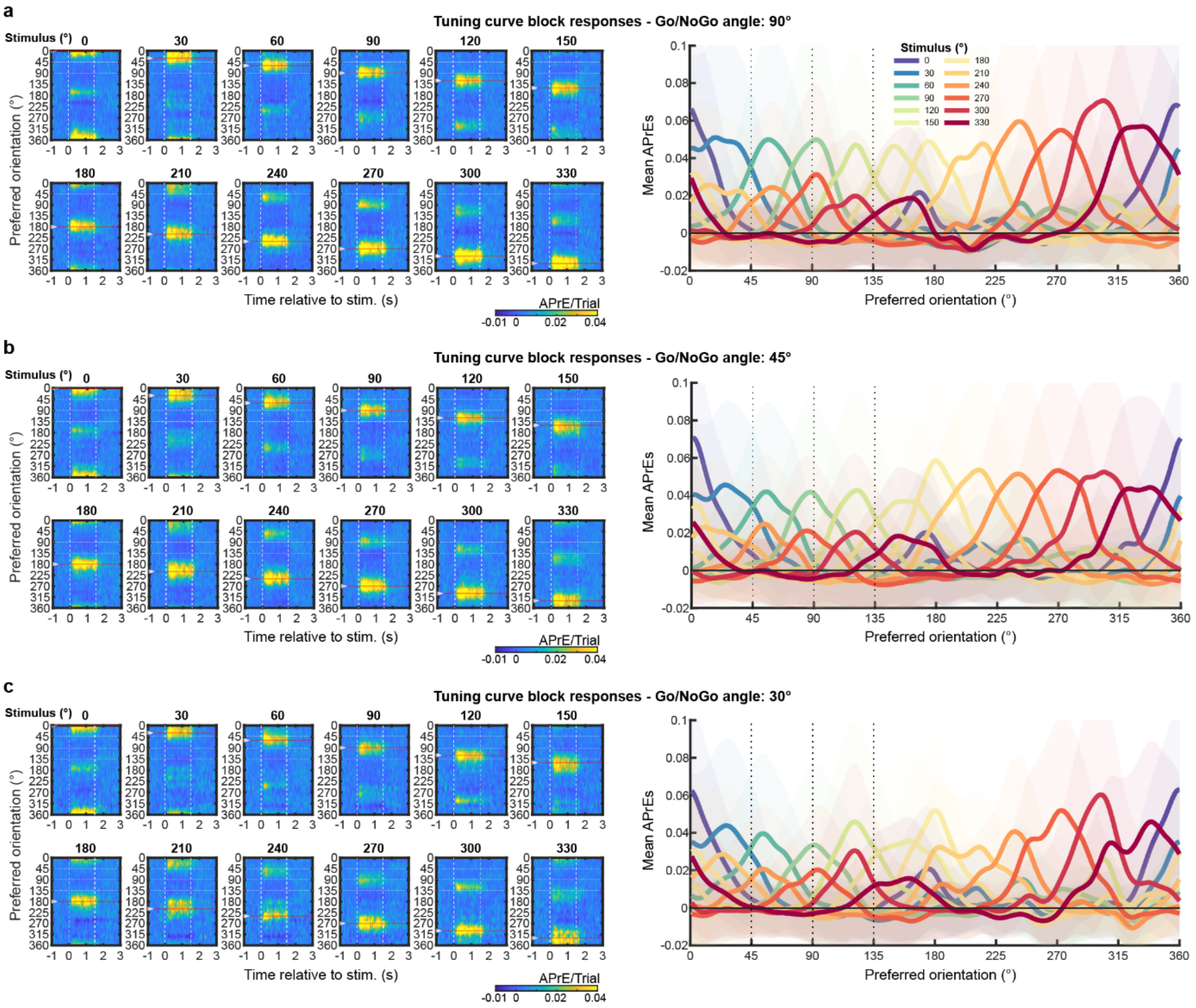

**Fig. S14: Representations in V1 of tuning block stimuli in naive animals between recording days 1-3. a.** *Left*: Neuronal activity evoked by the presentation of all 12 tuning block stimuli as a function of the neurons' preferred orientations. *Right*: Average activity profile in the preferred orientation space during the two first seconds of stimulus presentation for all stimuli of the tuning block following the recording session D1. Mean ± s.e.m. **b.** Same representation as in a. for the recording session D2. **c.** Same representation as in a. for the recording session D3.

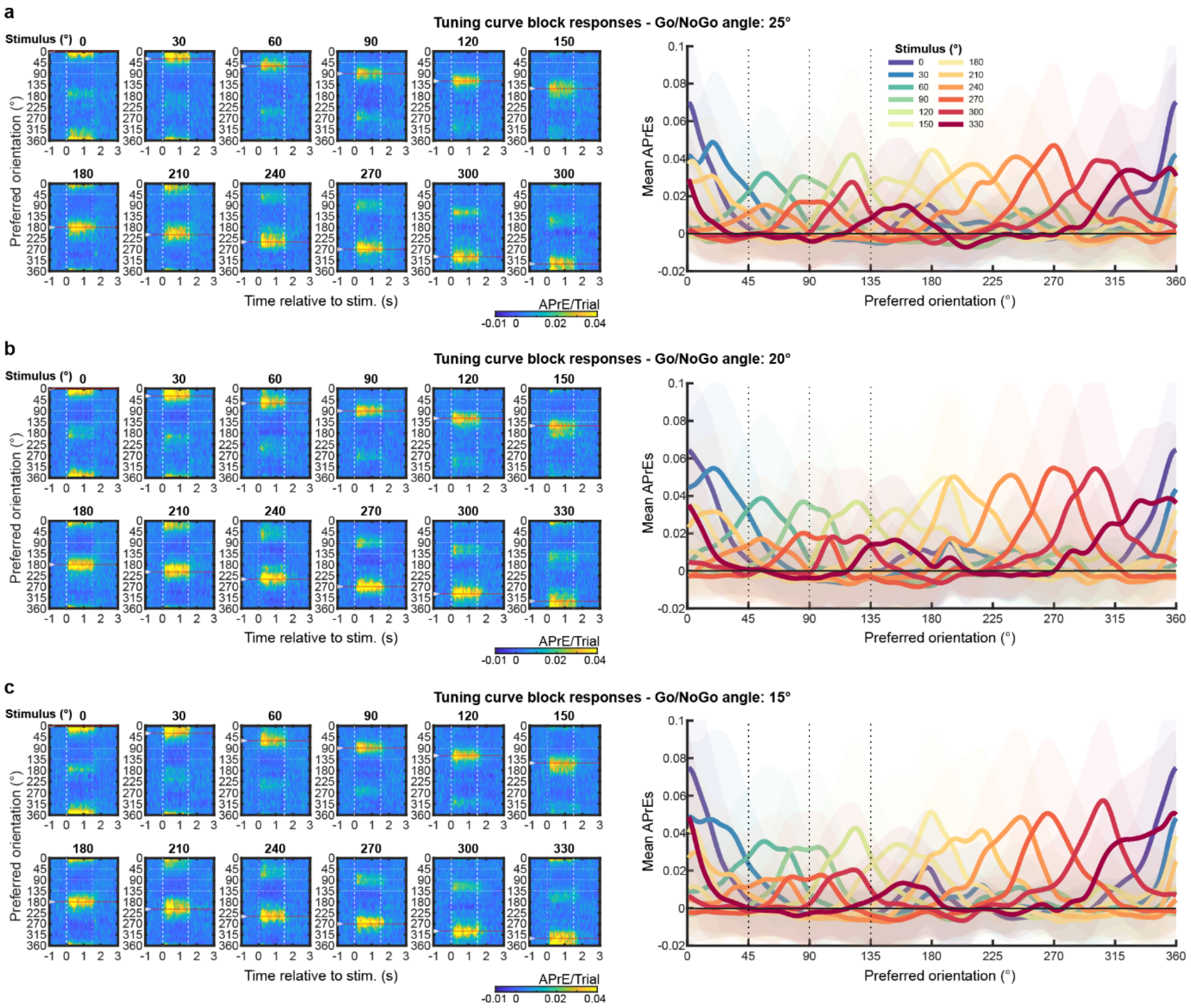


**Fig. S15: Representations in V1 of tuning block stimuli in naive animals between recording days 4-6. a.** *Left*: Neuronal activity evoked by the presentation of all 12 tuning block stimuli as a function of the neurons' preferred orientations. *Right*: Average activity profile in the preferred orientation space during the two first seconds of stimulus presentation for all stimuli of the tuning block following the recording session D4. Mean ± s.e.m. **b.** Same representation as in a. for the recording session D5. **c.** Same representation as in a. for the recording session D6.

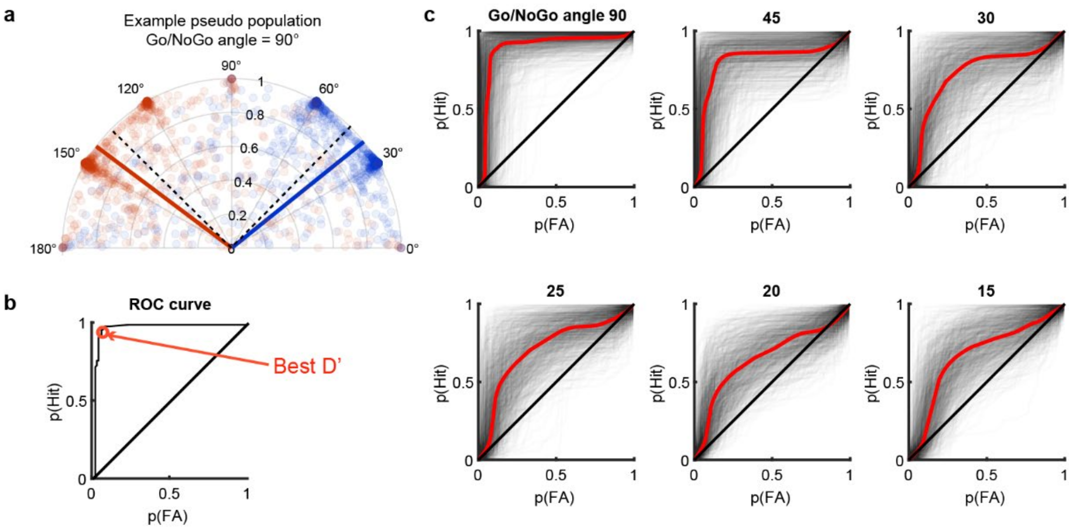


**Fig. S16: ROC analysis on the SNNs trained with the trials of the tuning block. a.** Example of the decision distribution for a SNN discriminating between the Go and NoGo cues on D2 when the angle between the task stimuli was 90°. This SNN was trained to discriminate between the 12 oriented stimuli of the tuning block. Each circle corresponds to the estimation for a single trial of the orientation of the task cues. The estimated orientation for a trial was defined as the semi-circular average of the SNN output vector for that trial. Go trials are labelled in blue and NoGo in red. The solid lines indicate the averaged orientation across all trials. The dashed lines indicate the actual stimuli orientations (45° and 90°). **b.** ROC curve obtained from the data in (a). The best D', corresponds to the point of the ROC curve that is the furthest from the bisector. It is indicated in red. **c.** All ROC curves (1000 per Go/NoGo angle), with the average curve across all Go/NoGo pairs in red.

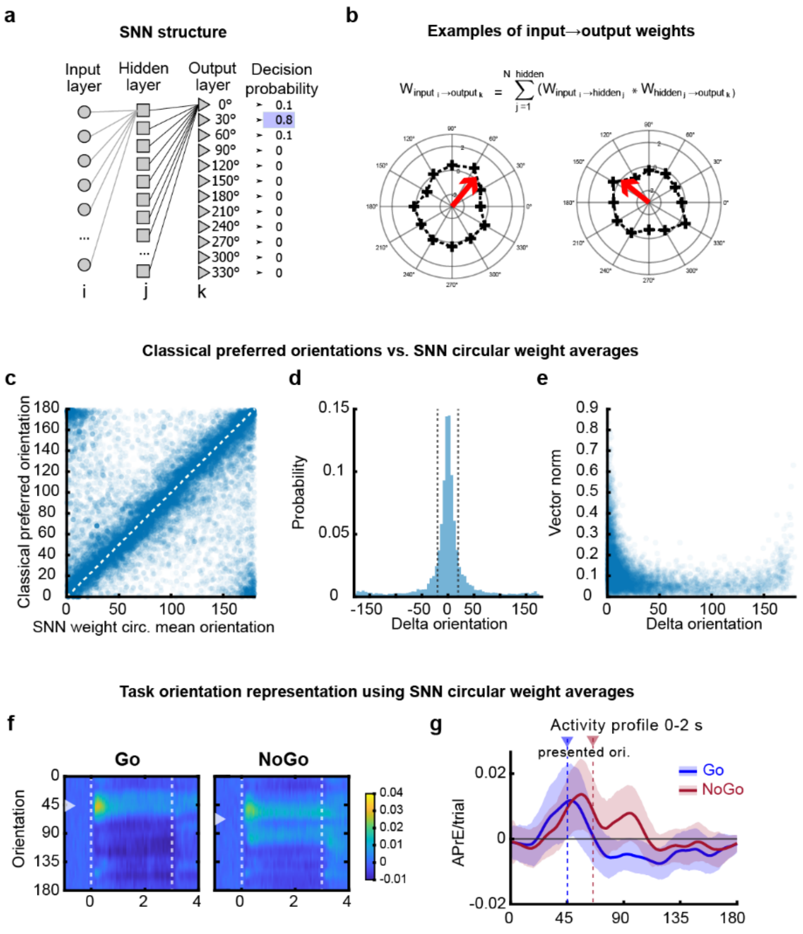


**Fig. S17: Orientation-trained SNNs use the same statistical properties of the neuronal responses that are captured by classical tuning curves. a.** Schematic of the SNN trained to classify stimuli from the tuning block. **b.** Examples of two sets of input→output weights, corresponding to the influence each neuron has on the decision output. For each input neuron, we can determine a twelve-weight vector that correspond to the respective weights this neuron has on each of the SNN output neurons (black crosses). The "Weight Orientation" (WO), the semi-circular average of the input neurons' weights, is indicated in red. **c.** Scatter plot of the SNN weight orientations vs. the classical preferred orientations (PO) of all SSN input neurons. Most of the values are similar and sit on the bisector **d.** Distribution of the differences between SNN WO and PO. The median absolute difference was 10.7°. The vertical dashed lines indicate 0° ± 20°; 67.2% of neurons are within these values. **e.** Norm of the vectors against the WO/PO difference. Neurons with a more selective weight distribution (resulting in a higher WO norm) have closer WO/PO values (25/50/75% quartiles of the delta orientation for neurons with the top half norms: 2.6/6.2/13.1°). **f.** Neuronal activity in the orientation space based on the WO values for the Go and NoGo stimuli (Go/NoGo angle 20°). **g.** Activity profile averaged over two seconds of stimulus presentation, from the data in (f). This profile is to be compared with the one obtained with our tuning curve fitting protocol to determine the preferred orientation and used in Fig 2.

| Dataset | Go/NoGo ang. | N animals Ca imaging | N animals behavior | N cells |
|---|---|---|---|---|
| Visual Only | 90° | 10 | 10 | 2007 |
| | 45° | 10 | 10 | 2524 |
| | 30° | 10 | 10 | 2600 |
| | 25° | 10 | 10 | 2627 |
| | 20° | 10 | 10 | 2594 |
| | 15° | 10 | 10 | 2675 |
| Audiovisual | 45° | 6 | 7 | 517 |
| | 30° | 7 | 10 | 779 |
| | 25° | 6 | 7 | 565 |
| | 20° | 4 | 7 | 290 |
| | 15° | 6 | 9 | 552 |
| Naive | 90° | 7 | - | 13252 |
| | 45° | 7 | - | 13455 |
| | 30° | 7 | - | 11542 |
| | 25° | 7 | - | 12065 |
| | 20° | 7 | - | 12303 |
| | 15° | 7 | - | 12657 |

**Table S1. Datasets description.**
